# Connecting polygenic disease risk to cell states and regulatory programs through single-cell chromatin accessibility

**DOI:** 10.64898/2026.04.27.721080

**Authors:** Liyang Yu, Luke T. Deary, Qiaoxue Liu, Qirui Zhang, Siming Zhao

## Abstract

Single-cell ATAC-sequencing enables high-resolution mapping of cell states and their putative regulatory elements. The enrichment of genome-wide association study (GWAS) signals within context-specific regulatory regions can reveal disease-relevant cell populations. We present Single-Cell ATAC-seq Disease Score (SCADS), a computational framework that integrates GWAS summary statistics with scATAC-seq data to prioritize disease-relevant cells. SCADS produces calibrated, cell-level scores that are comparable across datasets and traits by: (i) identifying chromatin co-regulatory regions via topic modeling, (ii) quantifying their polygenic GWAS enrichment, and (iii) calculating cell-specific disease scores from topic weights and enrichment levels. Across extensive simulations, SCADS outperforms existing methods in power while controlling false positives. Applied to multiple autoimmune traits, SCADS reveals marked heterogeneity in disease relevance within canonical immune cell types. We specifically studied the differential disease relevance within CD8^+^ T and colon epithelial cells for inflammatory bowel disease, and pinpointed underlying gene programs and variants. SCADS is scalable, interpretable, and modular, providing a general framework to connect noncoding regulatory variation to cellular identity and function.

## Introduction

Identifying the tissues and cell types that contribute to a disease is a crucial step toward understanding disease mechanisms. Many complex diseases exhibit substantial heritability, indicating that genetic variants can provide valuable insights into disease etiology^1^. Genome-wide association studies (GWAS) have successfully pinpointed thousands of loci associated with disease ^2–4^. Because the majority of disease-associated variants lie in regulatory regions, which are often active in a cell-type or tissue-specific manner, integrating GWAS results with regulatory-element annotations offers a powerful means of delineating disease-relevant cellular contexts^5–7^.

Indeed, methods such as stratified linkage-disequilibrium score regression (S-LDSC)^8^ have been developed and applied to reveal disease-relevant cell types for a wide range of traits. Most current efforts infer disease-associated cell types using bulk ATAC-seq data or by constructing pseudo-bulk profiles for cell types identified from single-cell ATAC-seq (scATAC-seq)^5,9–18^; consequently, disease relevance is derived at the level of discrete cell types. However, cellular states often exist on a continuum rather than as strictly separate categories, leading to ambiguity in cell-type annotation and downstream analysis^19^. Moreover, analyses that treat each cell type as homogeneous ignore inherent heterogeneity within cell types and may therefore miss biologically important processes such as differentiation, activation, or environmental responses^20,21^.

The rapid accumulation of scATAC-seq data now provides unprecedented resolution of chromatin accessibility for individual cells^17,22–24^. Consequently, there exists great potential to map disease relevance to specific cellular states by integrating scATAC-seq with GWAS data. In principle, if we knew the exact accessibility status of every regulatory element in an individual cell, we could apply S-LDSC^8^—as is routinely done for bulk ATAC-seq—to assess disease relevance. Specifically, by treating the cell’s open regions as an annotation, S-LDSC could estimate the per-variant heritability enrichment for variants in open regions relative to closed regions, yielding a disease score for that cell (**Fig. 1a**). In practice, this is hampered by the extreme sparsity of scATAC-seq: A typical cell captures reads in only ∼3 % (1-10%) of the regions that are detectable in bulk^23,25^, leaving the majority of regulatory elements unobserved and limiting the power of enrichment tests.

**Fig. 1.**
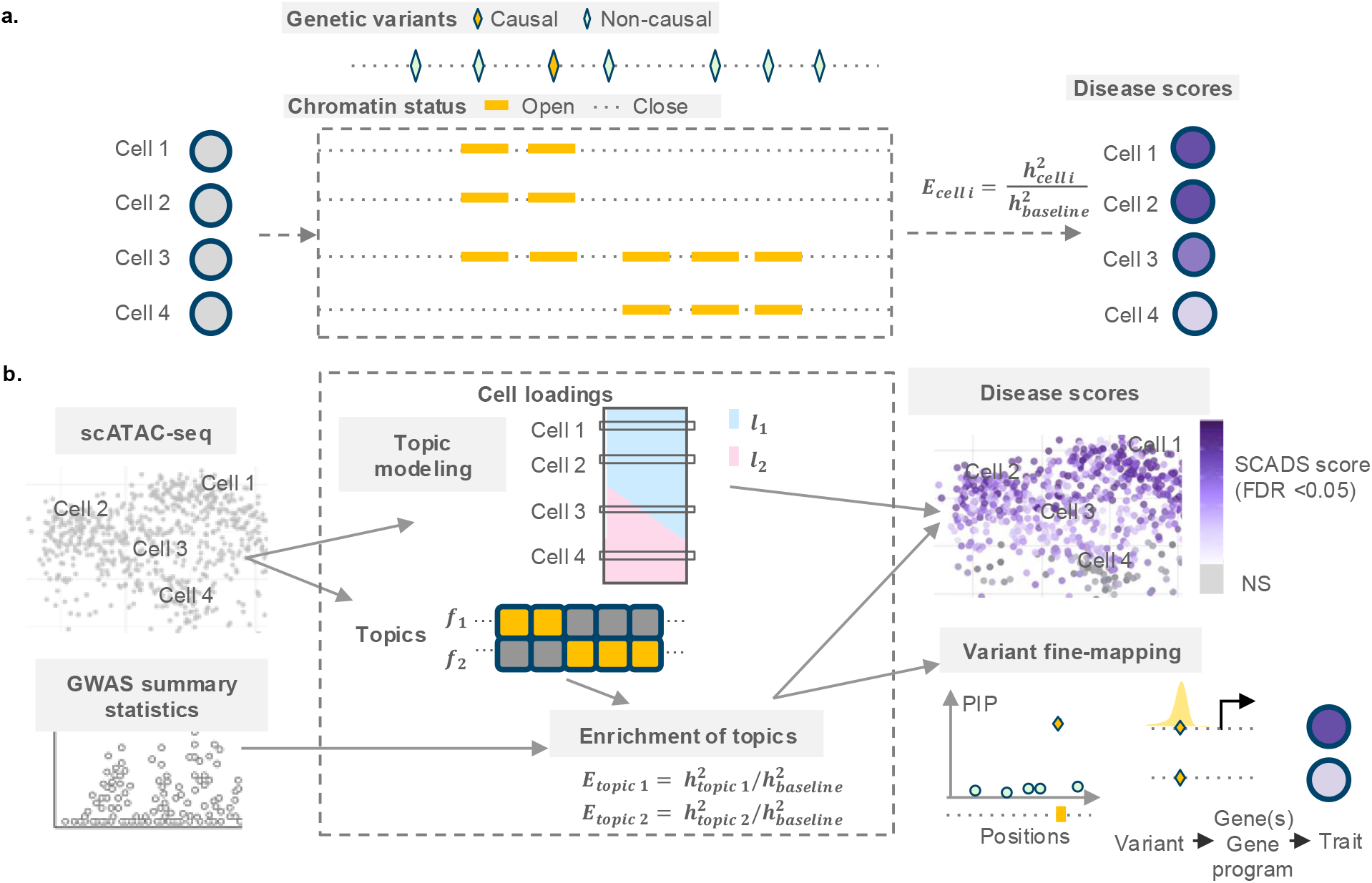
Overview of the SCADS method. **a**. When the chromatin statuses are known in each cell, an enrichment score for cell *i* (*E*_*cell i*_) is defined as the ratio of per-variant heritability in the open chromatin regions of the cell (*h*_*cell i*_^2^) to per-variant heritability genome-wide (*h*_*baseline*_^2^). **b**. The inputs for SCADS are the scATAC-seq count matrix and GWAS summary statistics. SCADS performs topic modeling on scATAC-seq data (UMAP of grey cells with four highlighted selections) to obtain (1) cell loadings (***l***_***k***_; topic *k*) and (2) topics (***f***_***k***_; topic *k*). The SCADS cell-level score, variance and *p* values are reconstructed from topic-level results and cell loadings (see Methods). On the UMAP on the right, darker colors indicate higher SCADS scores and more disease relevance, and non-significant (NS) enrichment is shown in grey. The outputs of SCADS assist with variant fine-mapping and more detailed study of variant to trait mechanisms.

To overcome this limitation, several recent methods have been proposed. SCAVENGE^26^ evaluates the enrichment of fine-mapped GWAS variants in open regions per cell but retains this enrichment as the disease relevance score only for the top-ranked cells (seed cells); the scores for the remaining cells are inferred from their similarity to these top cells. This strategy is constrained by the noisy, sparse information in individual cells and thus is prone to mistakes when choosing seed cells. It also does not exploit the polygenic architecture of most traits. scPRS computes a polygenic risk score for each cell based on read counts overlapping GWAS variants and then smooths the scores using a cell-cell similarity graph^27^. However, scPRS requires individual-level labeled training data, which may not be available for many traits. The scores produced by scPRS are also not directly comparable across traits or datasets and lack calibrated measures of uncertainty.

To address these shortcomings, we developed SCADS (Single-Cell ATAC-seq Disease Score), a method that infers disease relevance at the resolution of individual cells. The key idea is simple: We first project each cell onto a low-rank representation of its chromatin-accessibility profile, perform polygenic enrichment analysis in this reduced space using GWAS summary statistics, and then reconstruct a cell-level disease relevance score. It also simultaneously produces the set of regulatory regions driving that relevance which can be used to fine-map GWAS causal variants. Extensive simulations and applications to real data demonstrate that SCADS achieves higher power and lower false-positive rates than existing approaches across a variety of settings. Applying SCADS to a broad collection of scATAC-seq datasets and multiple complex traits, we pinpointed the cellular contexts most relevant to each trait and fine-mapped causal variants that are likely to act within those contexts, thereby delivering a more refined view of genotype-phenotype relationships.

## Results

### Overview of SCADS

SCADS provides a cell-level score that reflects a cell’s disease relevance, defined as the per-variant heritability in the open regions of the cell compared with background. The essential idea is that, although it is difficult to infer open chromatin regions from a single cell, learning the co-regulatory pattern of chromatin status across cells allows us to reliably estimate the contribution of a few “open-region sets” in any cell; we can therefore obtain the cell’s score by estimating the disease relevance of each “open-region set” and aggregate their contributions (**Fig. 1b**). More specifically, in the first step, SCADS performs topic modeling on scATAC-seq read-count data (see Methods); we employ the fastTopics^28,29^ implementation for computational efficiency. It then obtains topic loadings for each cell and defines an “open-region set” for each topic (see Methods), which are groups of co-regulated regions underlying a transcriptional program. In the second step, SCADS assesses disease relevance for each topic by calculating the heritability enrichment for a GWAS trait using S-LDSC, treating the corresponding “open-region set” as the functional annotation. Finally, SCADS constructs the cell-level score as the expected heritability enrichment for variants in the cell’s open regions, together with its variance and p-value (see Methods). The formula is intuitive given that the cell score depends on its topic loadings, the size of each topic, and the disease relevance of each topic. This workflow allows SCADS to overcome sparsity by projecting cells onto lower-dimensional factors, assessing disease relevance at the latent-factor level, and reconstructing the cell score. It leverages the polygenic trait signal for more powerful enrichment detection and provides a score with an intuitive meaning that is comparable across datasets.

SCADS further leverages the cell-level score and topic-derived functional annotations for downstream analysis (**Fig. 1b**). It uses the topic annotations and the learned enrichments as priors for Bayesian fine-mapping, and the cell scores enable novel investigations of variant-trait associations—for example, a positive correlation between the chromatin accessibility containing the variant and SCADS cell scores indicates that up-regulation of the variant’s target gene or a coregulated gene program is associated with the disease, thereby providing more mechanistic insights into variant-trait associations.

### Evaluating SCADS in simulations

We performed realistic simulations to benchmark SCADS against existing methods (see Methods). In brief, we simulated scATAC-seq data using the topic model. Topic-specific accessibility was derived from real scATAC-seq pseudobulk data of 5 blood cell types^30^. The total number of fragments per cell was sampled from a log-normal distribution matching empirical sequencing depth. We simulated GWAS traits using genotype data from 45K individuals from UK Biobank^31^, varying the fractions of causal variants, the causal variant enrichments in the 5 topics, and the total trait heritability corresponding to different test scenarios.

In the first set of experiments, we evaluated the performance of SCADS with different genetic architectures of the GWAS trait. In these simulations, a single topic had 20 fold (20x) higher enrichment of heritability while all other topics contained no enrichment. We set the total trait heritability to 0.5 but varied the number of causal variants. We showed results for two representative settings: One had ∼9300 causal variants (baseline fraction of causal variants, *π*^*b*^ = 1 × 10^−3^), (the dense setting, **Fig. 2a-c**), while the other had around 2300 variants (*π*^*b*^ = 2.5 × 10^−4^)(the sparse setting, **Fig. 2d-f**). We assessed the performance of SCADS and a comparator method, SCAVENGE, by comparing their cell scores with the true enrichment of heritability in the open regions of the cell. In the dense setting, SCADS correctly assigned the highest scores to the cells with true enrichment of heritability (comparing SCADS scores on the UMAP in **Fig. 2b** with the truth on UMAP in **Fig. 2a**), and the Spearman correlation between the SCADS score and the cell’s true enrichment was 0.85 (**Fig. 2b**). In contrast, SCAVENGE misidentified cells relevant to the traits (comparing trait relevant scores or TRS in **Fig. 2c** with **Fig. 2a**), and the correlation was 0.17. In the sparse setting, SCADS scores maintained high correlation with the true enrichment values (Spearman *r* = 0.77). While SCAVENGE improved its identification of the cells with true enrichment, the correlation was much lower (0.26) (**Fig. 2d-f**). Across all evaluated settings, we noted a consistent performance trend. SCADS remained robust across diverse genetic architectures, whereas the performance of SCAVENGE declined in settings with higher polygenicity (Supplementary Fig. 1j-l). This is expected as SCAVENGE relies on fine-mapped variants, which is sensitive to GWAS power, whereas SCADS utilizes S-LDSC to capture more polygenic signals.

**Fig. 2.**
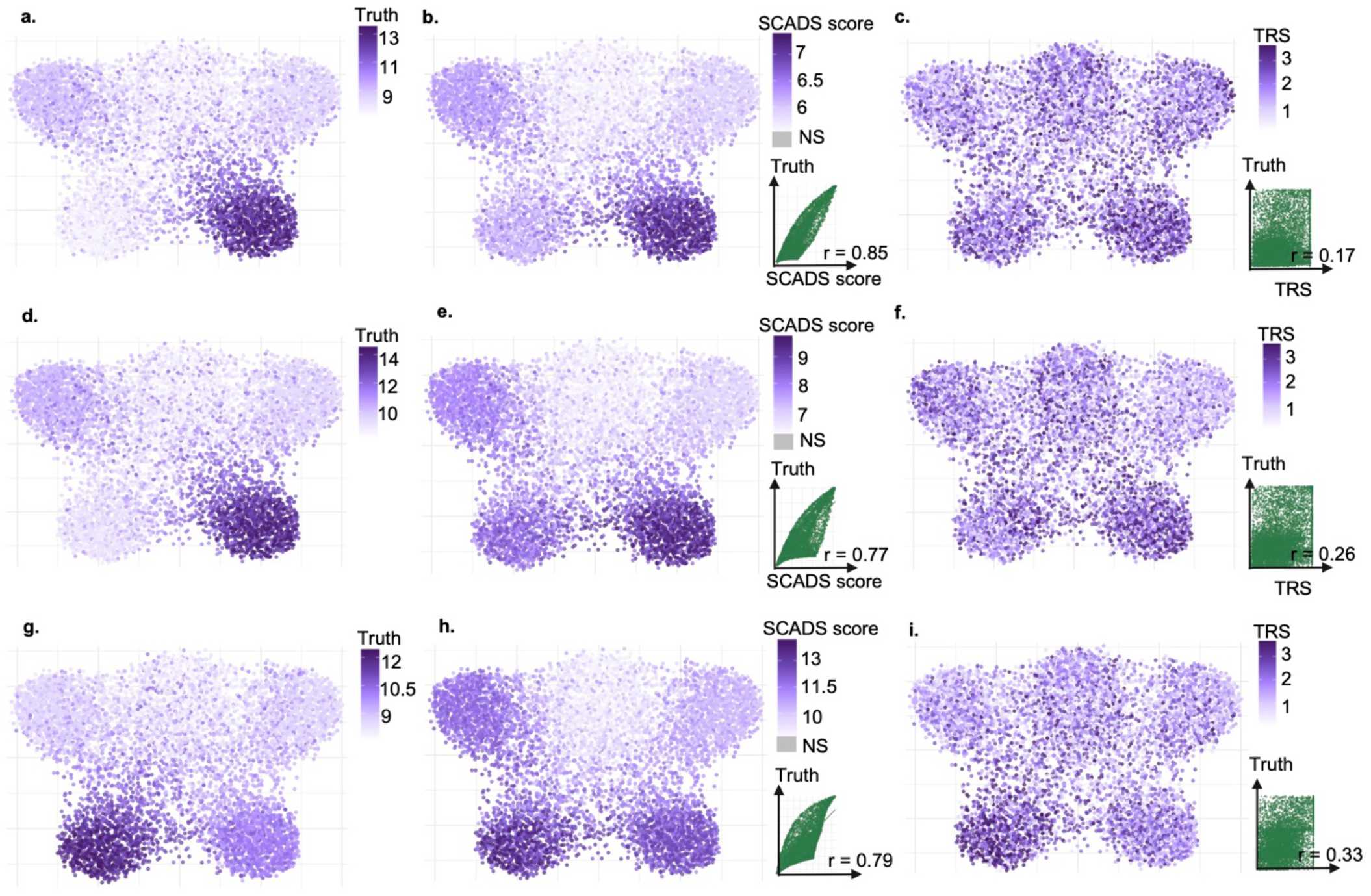
Evaluating the performance of SCADS in simulations. The simulated scATAC-seq data comprise 260K peaks across 7.5K cells (see Methods). **a-c**. Results for simulated GWAS with baseline fraction of causal variants *π*_*b*_ = 0.001 and one causal topic (fraction of causal variant in the casual topic open regions is 20x of *π*_*b*_). We calculated a true score for each cell, which is the true enrichment of heritability in the cell’s open regions. This is computed via Equation (1) using the known open chromatin status of each topic (*I*_*ik*_), the enrichment of heritability in topic open regions (*e*_*k*_) and the cell loadings (*l*_*ik*_) set in the simulation. UMAPs of cells are colored based on true score (**a**), SCADS score (**b**) and SCAVENGE TRS scores (**c**). **b-c**. Scatterplots of the predicted score (x-axis) vs. truth (y-axis) are shown on the lower right. Lines in these plots represent y = x. **d-f**. Results for simulated GWAS with baseline fraction of causal variants *π*_*b*_ = 2.5 × 10^−4^ and one causal topic (fraction of causal variant in the casual topic open regions is 20x of *π*_*b*_). **g-i**. Results for simulated GWAS with baseline fraction of causal variants *π*_*b*_ = 2.5 × 10^−4^ and two causal topics (fraction of causal variant in the casual topic open regions is 10x of *π*_*b*_ for each). For **d-f**. and **g-i**., the layout mirrors panels **a-c**. SCADS scores with FDR > 0.05 (non-significant or NS) are colored in grey.

Given that complex traits typically involve multiple cellular contexts, we moved on to a more complicated scenario, where we simulated GWAS data with enriched heritability (10x) in the accessible regions of two topics. We benchmarked using the sparse setting (*π*^*b*^ = 2.5 × 10^−4^) (**Fig. 2g-i**). On the UMAP of true cell enrichment values (**Fig. 2g**), we observed two cell clusters with enrichment, both at the bottom. SCADS highlighted these two clusters as well and exhibited a high correlation with the true values (Spearman *r* = 0.79) (**Fig. 2h**). In contrast, SCAVENGE only roughly recovered one cluster, resulting in a low correlation with the truth (Spearman *r* = 0.33) (**Fig. 2i**). Increasing the enrichment level in the causal topics (20x) improved SCAVENGE performance (Spearman *r* = 0.42), but still underperformed SCADS (Spearman *r* = 0.88) (Supplementary Fig. 1g-i).

Lastly, we performed null simulations with no true enrichment of heritability in any cells and evaluated SCADS’s control of false discovery rate (See Methods). We determined the *p* values from SCADS are well calibrated with slight deflation (more conservative) at the upper tail. (Supplementary Fig. 2a). Consistent with this, we detected a good control of false positives of SCADS in these simulations (Supplementary Fig. 2b).

### SCADS has high precision in identifying trait related cells

We next benchmarked SCADS performance using real GWAS data. We used monocyte count GWAS and evaluated if our method could recover monocytes as the most relevant context. In the first setting, we simulated scATAC-seq data from a pseudobulk scATAC-seq data for 2 cell types, B cells and monocytes, using an approach adopted from SCAVENGE (see Methods). We benchmarked SCADS with SCAVENGE and g-chromVAR^32^, a commonly used tool to assess enrichment of disease variants in accessible regions for scATAC-seq data. SCADS identified all monocytes as the cells associated with the monocyte count trait (100%), and the performance of SCAVENGE was comparable (98.6%), but g-chromVAR failed to identify ∼20% of monocytes (**Fig. 3a-c**).

**Fig. 3.**
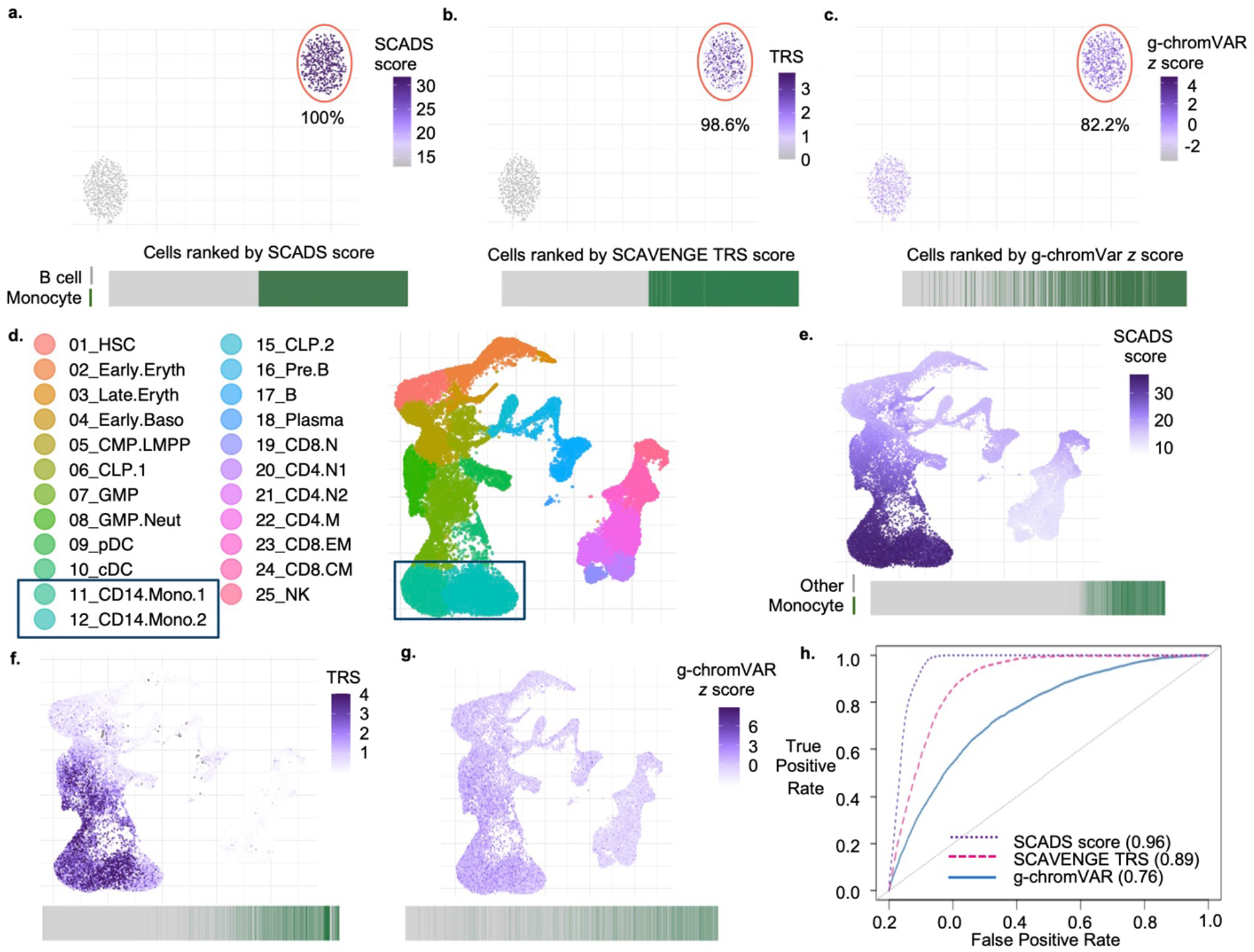
Benchmarking SCADS using monocyte count trait. **a-c**. Results using simulated scATAC-seq data with equal numbers of B cells and monocytes. UMAP plots of cells colored by SCADS scores (**a**), SCAVENGE TRS scores (**b**), and g-chromVAR z-scores (**c**) are depicted. For each method, we label the cells predicted as monocytes when the cell is ranked in the top half of scores. Monocytes are encircled in red on UMAP, and the percentage of true monocytes recovered by each method is noted below. Beneath each UMAP, a line plot ranks all cells from low to high score (left-to-right), with dark green lines representing true monocytes and gray lines representing true B cells. **d**. UMAP plot of the published hematopoiesis scATAC-seq data^18^ with monocytes highlighted in a black box on the legend and on the UMAP. CLP, common lymphoid progenitor; GMP, granulocyte-macrophage progenitor; MPP, multipotent progenitor; CMP, common myeloid progenitor; LMPP, lymphoid-primed multipotent progenitor; MEP, megakaryocyte-erythroid progenitor; BMP, basophil-mast cell progenitor; N.CD4 T, naive CD4^+^ T cell; N.CD8 T, naive CD8^+^ T cell; M.CD4 T, memory CD4^+^ T cell; CM.CD8 T, CD8^+^ central memory T cell; CD8.EM T, CD8^+^ effector memory T cell; Mega, megakaryocyte; Ery, erythrocyte; Baso, basophil; Mono, monocyte; Neut, neutrophil; cDC, classical dendritic cell. **e-g**. UMAP plots of the same cells as in **d**, colored by SCADS, SCAVENGE, g-chromVAR scores for the monocyte count trait. The gray lines in the line plot now denotes all other cells excluding monocytes. **h**. Receiver Operating Characteristic (ROC) curves are shown for each method with Area Under ROC (AUROC) values listed in parentheses.

In the second setting, we again assessed the monocyte count trait, but this time using the scATAC-seq data of 33,819 cells from human hematopoiesis, labeled with 23 known cell types^18^ (**Fig. 3d**). We visualized the cell scores from SCADS, SCAVENGE and g-chromVAR on UMAP (**Fig. 3e-g**). SCADS identified monocytes as having the highest scores, consistent with the expected biology of the trait. Then for each method, we ranked the cells from lowest to highest based on corresponding cell scores and obtained the AUROC values employing the cells with the monocyte labels as the ground truth. SCADS had the highest AUROC of 0.96, outperforming SCAVENGE (0.89) and g-chromVAR (0.76) **(Fig. 3h**).

### SCADS assesses disease relevance for several autoimmune traits at cell level

We applied SCADS on the hematopoiesis scATAC-seq data^18^ for eight autoimmune traits: Rheumatoid arthritis (RA), systemic lupus erythematosus (SLE), hypothyroidism (HTD), inflammatory bowel disease (IBD), primary biliary cholangitis (PBC), type-1 diabetes (T1D), atopic dermatitis (eczema), and asthma. We first computed the mean SCADS scores for each cell type (**Fig. 4a**), and the results were consistent with the cell type relevance findings for these autoimmune traits as previously described^9^. We then visualized the heterogeneity of trait relevance by calculating the difference between the 95th and 5th percentile (or range) of SCADS score for each cell type (**Fig. 4b**). B cells, T cells and NK cells were the more relevant cell types for most traits, but they also displayed substantial heterogeneity. We then examined the distribution of SCADS scores for all cells across cell types (**Fig. 4c**, Supplementary Fig. 3b-i). Along the canonical B cell developmental trajectory (HSC → CLP1 → CLP2 → pre-B → mature B)^33^, we observed a general increasing trend of SCADS scores for RA, SLE, HTD and PBC, suggesting that the B cell maturation is biologically relevant to the genetics of these traits^34^. Some pre-B cells exhibited SCADS scores that exceeded those of mature B cells for HTD and T1D, indicating that distinct cellular states within the pre-B stage may be particularly relevant to these traits (Supplementary Fig. 3j-k). For CD4^+^ T, CD8^+^ T and NK cells, there were no more refined cell type labels to delineate different stages of the cell, but the scores distributions were highly correlated for all traits, suggesting that there are common disease-relevant underlying processes.

**Fig. 4.**
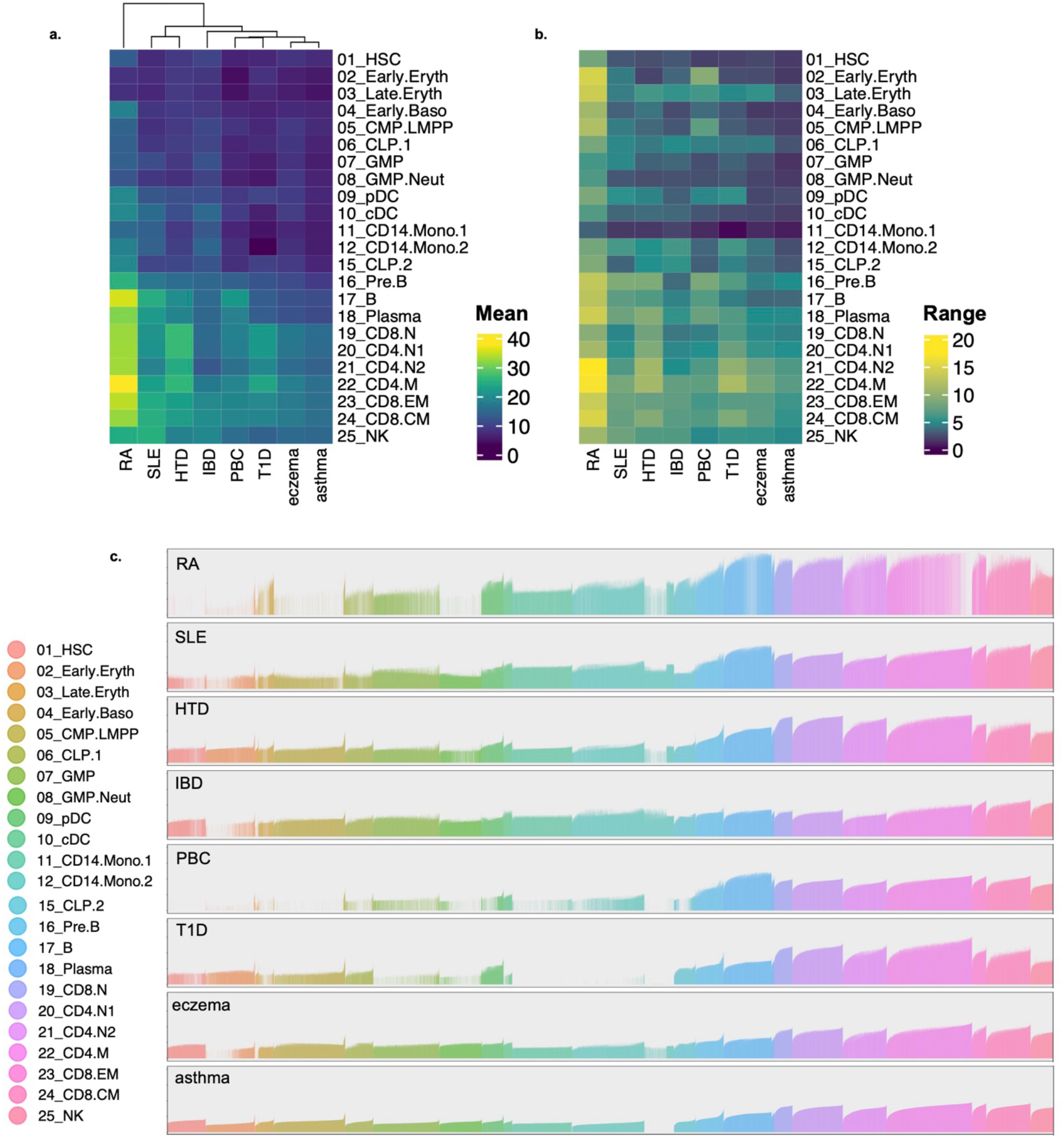
SCADS highlights disease relevant cell types and heterogeneity within cell types for various autoimmune traits. **a**. Heatmap of the mean of the SCADS scores for each cell type and trait. Cell type labels were obtained from hematopoiesis data^18^ (same as in **Fig. 3d**). Hierarchical clustering was performed for traits (column-wise). **b**. Heatmap of the range of the SCADS scores for each cell type and trait. Range is defined as the difference between the 95th and 5th percentile of scores. The order of the traits and cell types is the same as in **a. c**. Bar plots of SCADS scores for all cells for each trait. Each bar represents a cell with the height of the bar specifying the SCADS score. The cells are ordered based on their cell type labels as in **a** and **b**. Within each cell type, cells are ordered based on ascending SCADS score for the asthma trait. Color indicates cell type. Cells with FDR > 0.05 have a bar height of 0.

### Probing into the heterogeneity of IBD relevance in CD8^+^ central memory (CM) T cells

We further investigated the observed heterogeneity of IBD relevance in CD8^+^ CM T cells (**Fig. 5a**). This cell population appeared to be one of the cell types with the highest SCADS scores, and the scores formed a prominent gradient within this cell type. To study the mechanisms underlying this gradient, we correlated the topic loadings with the SCADS score (**Fig. 5b**). A positive correlation indicates that regulatory programs associated with the topic are likely to be disease relevant. We analyzed the topic with the highest positive correlation (Pearson *r* = 0.60), topic 8, which also had the highest enrichment of heritability (*e* = 21.23, *p* = 1.11 × 10^−5^). Transcription factor (TF) motif enrichment analysis of topic 8 revealed that the binding motifs of *ETS1* and ETS family TFs were among the top enriched motifs (See Methods; Supplementary Table 1). *ETS1* is pivotal in CD8^+^ CM T cells, regulating lineage commitment and effector function^35^. Dysregulation of this TFs network has direct implications for IBD pathogenesis, as ETS1 has been shown to facilitate Th1 cell-mediated mucosal inflammation through CIRBP upregulation^36^. Other top ranked TFs included RUNX, IRF1, IRF2, etc., which are essential regulatory factors in immune responses, immune cell development and differentiation^37,38^. Complementing this analysis, we compared cells with higher versus lower SCADS scores and identified differentially accessible chromatin regions (see Methods). Regions with increased accessibility in the higher-scoring group were enriched for binding motifs of RUNX1, RUNX2 and ETS family TFs (Supplementary Table 2), whereas regions with decreased accessibility were enriched for motifs of CTCF and BORIS (Supplementary Table 3). Together, these analyses implicate the ETS and RUNX transcription factor families as potential key players of IBD-relevant gene regulatory programs.

**Fig. 5.**
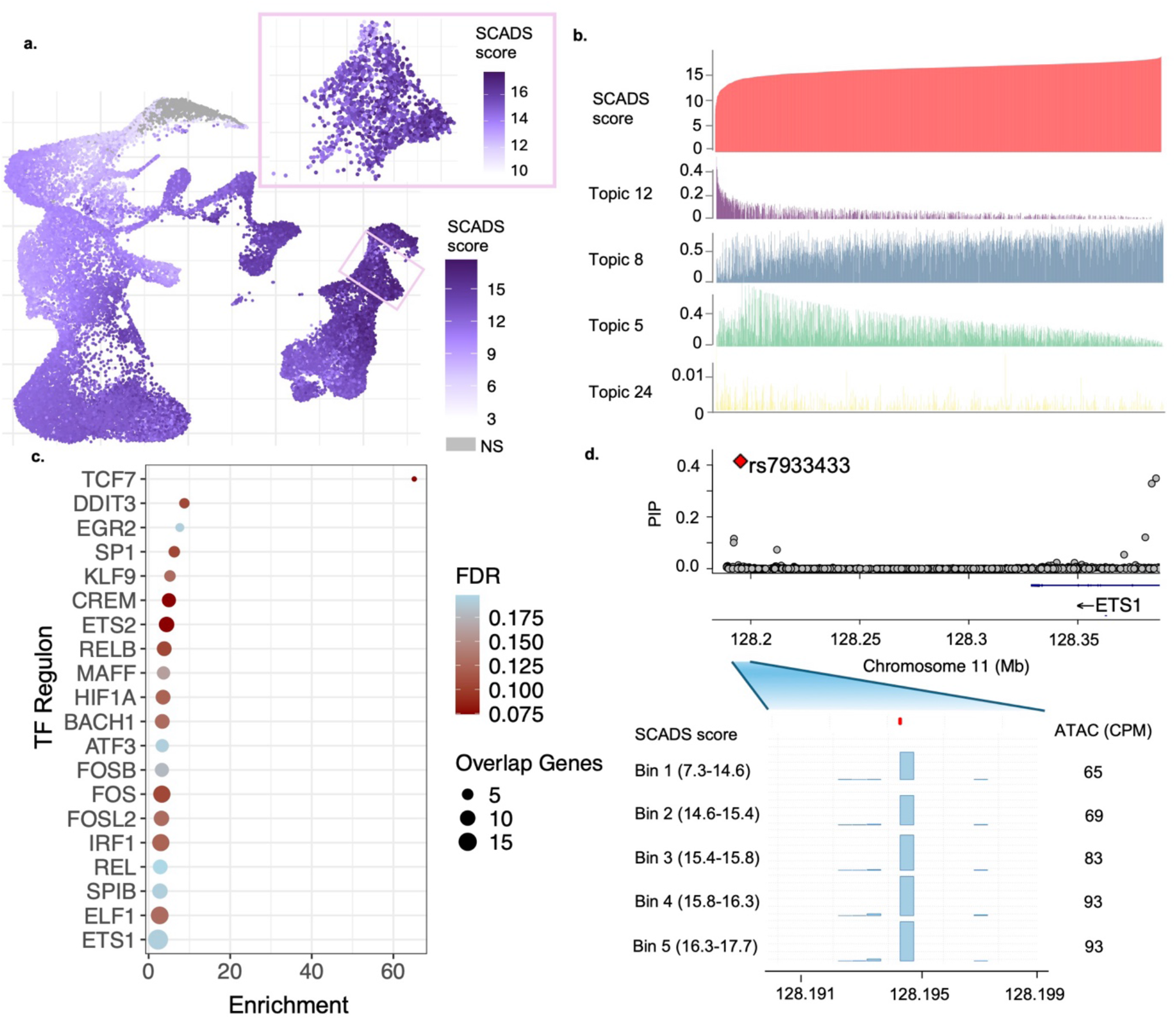
SCADS prioritizes CD8^+^ CM T cells specific regulatory variant(s) and mechanisms for IBD. **a**. UMAP plot of scATAC-seq hematopoiesis data of IBD SCADS scores, where the CD8^+^ CM T cells are featured in the pink box. **b**. Bar plot of SCADS scores and topic loadings for CD8^+^ CM T cells. Cells are ordered by SCADS scores from low to high. Corresponding cell loadings for each of the top four correlated topics (Pearson |r| > 0.2) are plotted below. **c**. Enrichment of IBD GWAS genes in TF regulons in immune cells. An IBD GWAS gene is assigned as the nearest gene to a variant with PIP > 0.3 and |cor| > 0.2. A Fisher’s test is then performed for each regulon. BH procedure is applied to get FDR (See Methods). The plot depicts top significant regulons (FDR < 0.2). **d**. Fine-mapping of ‘rs7933433’. In the top panel, PIPs for variants in the rs7933433 region (chr11:128.18Mb-128.38Mb) is shown. The bar plot below displays the scATAC-seq signals for cells grouped into 5 quantile bins based on their SCADS scores. The score ranges for each bin are on the left. The scATAC-seq signal is measured as counts per million (CPM), which is labeled on the right.

We next performed functionally informed fine-mapping of IBD GWAS variants employing topic annotations (see Methods). We identified 177 variants with Posterior Inclusion Probability (PIP) > 0.3. Of these, we further examined 35 variants whose chromatin accessibility correlated with the SCADS scores (see Methods) (Supplementary Table 4). We reasoned that the effect of such variants may aggregate at the gene program level; those whose accessibility positively correlates with the disease score may regulate predisposing genes, while those with the negative correlation may regulate protective genes for IBD. Thus, we compiled these variants and tested if the target genes of these variants aggregate in any regulatory programs. For this analysis, we first assigned the nearest protein-coding gene to each variant as its putative target gene. Then we identified the regulons, where each regulon included a TF and its set of transcriptionally co-regulated target genes, by applying SCENIC^39^ on a single cell RNA sequencing (scRNA-seq) data of immune cells^40^. Lastly, we inspected if any regulons were enriched for our set of target genes (see Methods).

We identified 20 regulons significantly enriched with our target genes (FDR < 0.2) (**Fig. 5c**, Supplementary Table 5). The most enriched regulon was the TCF7 regulon. TCF7 is crucial for differentiation and maintenance of CD8^+^ CM T cells, which are immune cells that provide long term protection against recurring threats^41,42^. One of the genes in the TCF7 regulon is the *ETS1* gene, whose binding motif we found to be highly enriched in the most disease-relevant topic (Supplementary Table 1). Of the 35 causal variants we had examined, rs7933433 had the second strongest positive correlation (Pearson *r* = 0.964) (**Fig. 5d**) between accessibility and the disease score. This variant is located in a multi-enhancer hub for *ETS1* and controls its dosage in T cells^43–45^. This establishes *ETS1* as a susceptibility gene whose dysregulation contributes to IBD pathogenesis, potentially by interacting with the TCF7 transcriptional regulatory network.

Beyond TCF7, ETS2 emerged as a significant regulon in our analysis (FDR < 0.1). Functional genomic studies have demonstrated that an intergenic region on chr21q22, independently linked to IBD, ankylosing spondylitis, and primary sclerosing cholangitis, increases *ETS2* expression specifically in macrophages, driving inflammatory gene programs^46^. This finding had been validated using single-cell colocalization approaches, which identified an *ETS2* eQTL colocalization with IBD GWAS signals in non-classical monocytes (CD16+)^47^. The ETS2 regulon includes several well-known IBD-relevant target genes such as *IRF1*^48^, *IL10*^49^, and *PTPN2*, the latter being a well-established IBD susceptibility gene that regulates T cell differentiation and intestinal barrier function^50,51^. We suspect the ETS2 regulatory network contributes to the disease relevance of these CD8^+^ CM T cells in IBD.

### SCADS reveals heterogeneity in colon epithelial cells for IBD relevance

Besides immune cells, the colon epithelium has also been shown to be an important player in IBD ^52,53^. Therefore we applied SCADS on a scATAC-seq data of colon epithelium^54^. In the UMAP of this scATAC-seq data (**Fig. 6a**), the larger cluster observed on the right comprises cells at various stages of enterocyte differentiation, spanning from stem cells to actively cycling transit-amplifying (TA) cells to premature and terminally differentiated mature enterocytes. This continuum formed a smooth differentiation trajectory (Supplementary Fig. 4). Intriguingly, the SCADS scores aligned precisely with this trajectory, displaying a clear gradient: Lower scores were observed in stem-like cells while higher scores reflected the more differentiated cell types (**Fig. 6b**). This implies that enterocyte differentiation constitutes a pivotal process in the context of IBD pathogenesis, and pinpoints mature enterocytes as the most disease relevant cells.

**Fig. 6.**
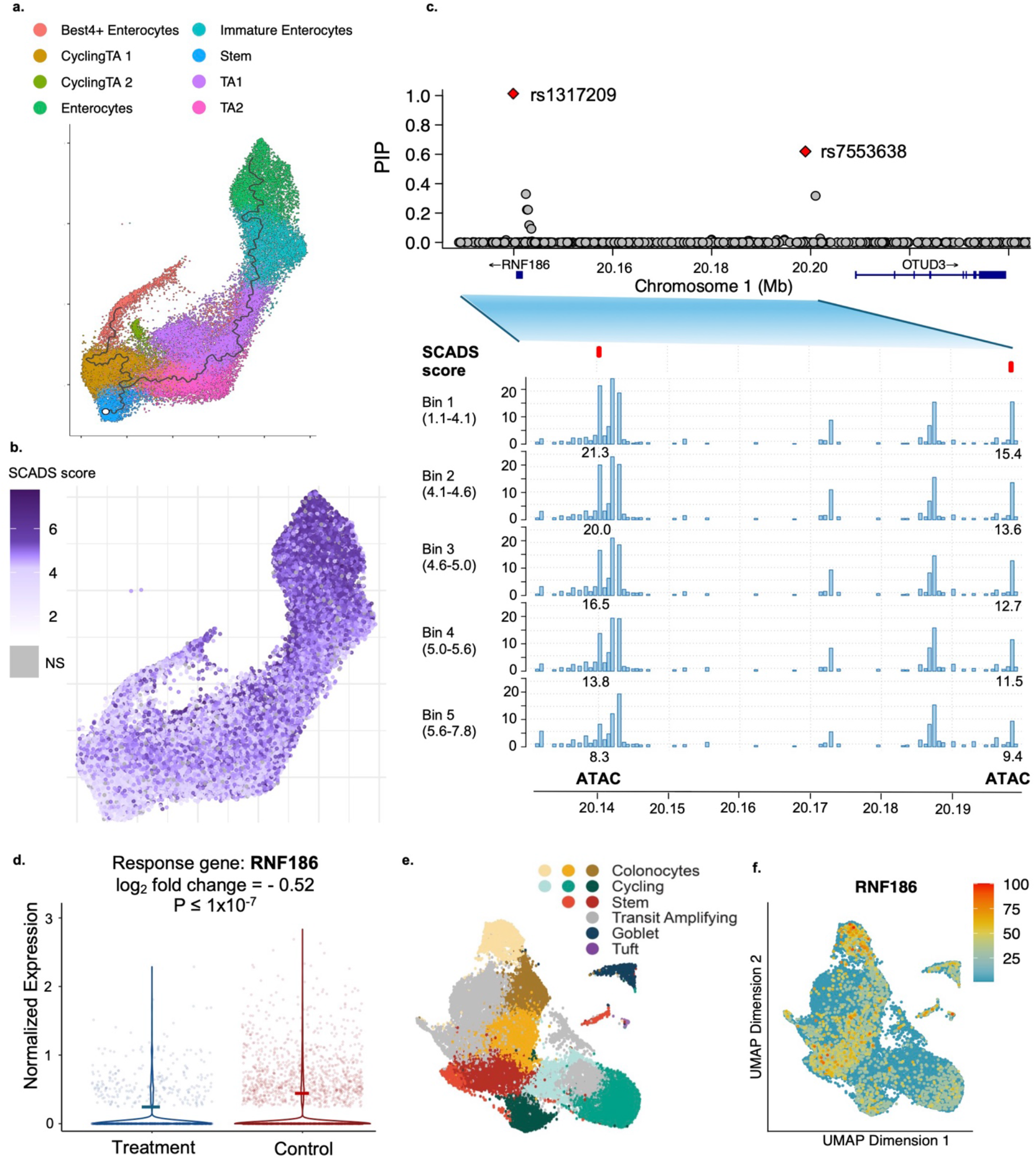
SCADS identifies colon epithelial cells–specific functional regulatory variant(s) linked to IBD. **a**. UMAP plot of cell types in colon epithelial cells of the intestine tissue^54^. Cell type labels were obtained from the same publication. The ragged black line on the right corresponds to the differentiation trajectory based on pseudotime (Supplementary Figure 5). **b**. UMAP plot of IBD SCADS scores for the colon epithelial cells. **c**. PIP plot for the genomic region around rs1317209 and rs7553638. The bar plot below shows the CPM for cells grouped into 5 bins based on their SCADS scores for a zoomed-in region (chr1: 20.13Mb-20.20Mb). **d**. Differential expression of *RNF186* from targeting rs7553638 with CRISPRi (Treatment) versus non-targeting (Control) in a colon organoid sample. **e-f**. UMAP plots of scRNA-seq data from the colon organoid. **e**. Cell types annotated based on known epithelial cell type markers (see Methods). **f**. *RNF186* gene expression across single cells from the CRISPRi experiment, which is normalized via SCTransform and displayed as relative expression intensity (scale of 0–100).

In contrast, we explored another trait, colorectal cancer (CRC), which also arises from dysregulation of the colon epithelium, but SCADS revealed that these diseases exhibit distinct cellular vulnerability profiles. Specifically, the stem cell compartment presented with the highest disease cell scores for CRC (Supplementary Fig. 5). This aligns with extensive evidence that LGR5+ intestinal stem cells serve as the cells-of-origin for CRC^55^, and that stemness-associated gene signatures not only identify cancer stem cells within tumors but also predict disease relapse^56^. These divergent scoring patterns underscore the biological distinction between IBD and CRC pathogenesis.

Returning to IBD, we then proceeded to fine-map variants leveraging topic derived annotations and identify those whose chromatin accessibility correlated with the SCADS score (see Methods). Again, we grouped variants into three groups: positive, negative, and no correlation (Supplementary Table 6). In the negative correlated group, we identified two top causal variants of significant interest (rs1317209 and rs7553638) (**Fig. 6c**). These two variants are spatially close (58.9kb apart) and display a co-accessibility pattern, suggesting that the encompassing genomic region is subject to a mechanism of coregulation. The accessibility decreased as the IBD SCADS score increased, suggesting a protective effect of their target genes. rs1317209 received a PIP of 0.999, and it has been speculated that *RNF186* is a potential target gene for rs1317209 due to its proximity. Its accessibility correlation with SCADS score was -0.978 (Pearson r). rs7553638 had a PIP of 0.619 and its causal variant role is less definitive, but its accessibility correlation with SCADS score is stronger (Pearson *r* = -0.990). The gene nearest to this variant is *OTUD3*, with *RNF186* being the second closest.

To further investigate the inverse correlation between the accessibility of rs7553638 and the SCADS score, we performed experiments in a colon epithelial organoid model to identify its target gene(s). Specifically, we utilized an organoid line derived from normal colon epithelium and employed CRISPR interference (CRISPRi) to experimentally target the rs7553638 locus (see Methods). We discovered it led to a significant reduction in *RNF186* gene expression (log_2_ fold change = -0.52, *p* = 1 × 10^−7^) (**Fig. 6d**) (see Methods). RNF186 encodes an E3 ubiquitin ligase, an enzyme that is critical for marking proteins for degradation or modification^57^. Its activity is crucial for maintaining the integrity and function of the intestinal mucosal barrier, which acts as the primary defense against gut microbes^58^. This is consistent with our expectation that the target gene of rs7553638 has a protective function in IBD. Using scRNA-seq data from the same organoid line, we found *RNF186* is highly expressed in colon epithelial cells, particularly in stem cells and colonocytes (**Fig. 6e-f**). These analyses support the role of rs7553638 as a causal variant and *RNF186* as a causal gene, also establishing enterocytes as the critical cellular context through which their effects on IBD are mediated.

## Discussion

We developed SCADS, a computational framework that prioritizes disease-relevant cell types and states by integrating GWAS summary statistics with scATAC-seq data. SCADS generates a cell-level score that reflects the enrichment of disease heritability within the cell’s open chromatin regions. Extensive simulations demonstrated that SCADS can accurately identify disease-relevant cells while effectively controlling for false positives. We demonstrated the utility of SCADS by applying it to autoimmune traits, particularly IBD, leading to novel insights into the disease’s genetic etiology.

The extreme sparsity of scATAC-seq data hampers the cell-level inference of disease relevance. Recent studies^59^ have shown that it is virtually impossible to determine whether a given chromatin region is open or closed in an individual cell. SCADS tackles this limitation by applying topic modeling to produce a denoised, covariate adjusted, low-dimensional representation for open or closed chromatin status and then performing cell-level inference from this low dimensional space. Other cell level inference methods also perform aggregation in some format to make cell-level inference. For example, SCAVENGE and g-chromVAR aggregate read counts across GWAS causal variants and compare against the background for a cell. scPRS uses Latent semantic indexing (LSI) to learn embeddings for individual cells to get a score. A few cell scoring methods for single-cell RNA-seq or multiome have been developed^60–63^, aggregating signals at disease gene sets or pathways. We found topic modeling is convenient for this application because it learns the “topics” from scATAC-seq data, which are often coregulated genomic regions with clear biological meanings. SCADS then reconstructs the cell-level score using the weights of the topics for a cell (determined by reads distribution and topic open region size) and the topic’s enrichment of GWAS signals compared to the background. The score has an explicit meaning–it is the enrichment of GWAS signals in the open regions of cells compared to the background.

We further demonstrate the utility of SCADS for uncovering disease-associated genetic mechanisms. By leveraging topic-derived annotations, we can infer gene-regulatory programs that drive disease susceptibility and perform fine-mapping to pinpoint causal variants. Exploiting the heterogeneity of CD8^+^ CM T cells, our analysis prioritizes several regulons that appear to underlie IBD. We further studied a casual variant whose openness correlates with the SCADS score and that acts on one of the regulons we identified. Using SCADS scores derived from colon epithelial cells, we pinpointed rs7553638 as a causal IBD variant despite its moderate PIP. Functional assays showed that this variant regulates *RNF186*, a gene with critical epithelial functions. Such analyses provide more mechanistic insights about the variant, its regulatory mediators, the cellular contexts and the disease.

A current limitation of SCADS is that it does not yet model batch effects or other covariates. Because scATAC-seq data remain scarce—particularly for difficult-to-obtain samples—there is an emerging need to integrate multiple experiments. This could be addressed by adopting more sophisticated topic-modeling frameworks (e.g., the GBCD model^64^) that explicitly account for batch effects. It is also possible to leverage mouse scATAC-seq datasets, which have proven to be valuable surrogates for human regulatory landscapes. We will therefore assess the utility of mouse scATAC-seq data for elucidating human traits in future studies.

In summary, we anticipate that SCADS will be broadly applicable to scATAC-seq profiles across diverse cell states and experimental conditions, thereby facilitating the understanding of the genetics of complex traits.

## Methods

### Topic modeling on scATAC-seq data

SCADS uses the fastTopics^29^ model to perform topic modeling for scATAC-seq read count data. In brief, we have

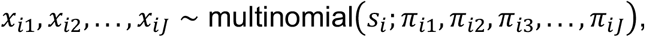

where *x*_*ij*_ is the single cell read count for cell *i* in region *j*. Assuming there are *K* topics, we have 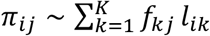, and 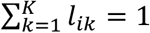. It assumes a region has a grade of membership *l*_*ik*_ for topic *k*. Let *I*_*ij*_ be an indicator of whether chromatin is open or not in cell *i* for region *j*. We model it as *I*_*ij*_ ∼ Bernoulli 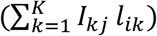, where *I*_*kj*_ represents if the region *j* is open or not in topic *k*.

In practice, we apply topic modeling to get *f*_*kj*_ and *l*_*ik*_ from the scATAC-seq read count matrix (cell by peak) and then infer *I*_*kj*_ based on this result (see below). For the rest of the genome (outside the peaks in the count matrix), we assume *I*_*kj*_ = 0.

### Deriving topic annotations for S-LDSC

To infer the topic-level open or close status *I*_*kj*_, we estimate the probability of *f*_*kj*_ > *f*_*kj*0_, where *f*_*kj*0_ is the baseline rate for region *j* in topic *k*. First, we estimate *f*_*kj*0_ accounting for GC count of region *j*. For bulk ATAC-seq data, a mixture model that treats read counts as a combination of baseline and true signals (gcapc^65^) has been developed. Under this model, it will give a GC-content-dependent bias curve for the baseline rate. SCADS generates a pseudobulk read count for topic *k*, region *j* as 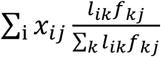; it then applies the gcapc model on the topic *k* pseudobulk data to predict a baseline rate based on the GC content of the region. This baseline rate is further normalized by the total reads for the topic ∑_*j*_ ∑_*i*_ *l*_*ik*_ *x*_*ij*_ to get *f*_*kj*0_. Next, we obtain the maximum a posteriori estimation of the log_2_ fold change (LFC), log_2_(*f*_*kj*_/*f*_*jk*0_), and quantify its uncertainty using the same algorithm as described in fastTopics. We then calculate a *z* score and a one-sided *p* value based on the estimated LFC and its error. After which we apply the Benjamin-Hochberg (BH) procedure and assign the region as open (*I*_*kj*_ = 1) at false discovery rate (FDR) < 0.05. Lastly, we construct variant annotations for topic *k*, denoted as *A*_*k*_, based on ***I***_***k***_ = {*I*_*kj*_, *j* = 1, …, *J*}. In practice, we suggest the pseudobulk read counts for a topic to reach 10 million in order to be sufficiently powered to derive the annotation.

### Cell score and variance

Let *h*_*j*_ be the heritability and *v*_*j*_ be the number of variants in region *j*. When *I*_*ij*_ is observed for cell *i*, the cell-level enrichment is defined as

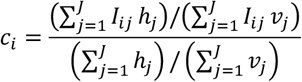

This means that we define the cell-level enrichment as the ratio of per variant heritability for variants in open chromatin regions of the cell to the genome-wide per variant heritability. This cell-level enrichment is the same as the enrichment of a functional annotation *A*_*i*_ defined by S-LDSC, when we form *A*_*i*_ for variants based on {*I*_*ij*_, *j* = 1, …, *J*} . Let *τ*_0_ be the genome-wide per variant heritability, so we have 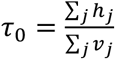, and *τ*_*i*_ be the per-variant heritability for variants in open chromatin regions in cell *i*, 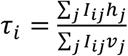. Hence, *c*_*i*_ = τ_*i*_ /*τ*_0_ . Although *I*_*ij*_ is unknown from scATAC-seq directly, we can model it as *I*_*ij*_ ∼ Bernoull*i* 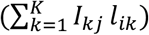 given the topic-level open or close status (*I*_*kj*_) and “grade of membership” (*l*_*ik*_) from the topic modeling of the scATAC-seq data. We use the expected value of *I*_*ij*_ to replace *I*_*ij*_ in the cell-level enrichment definition, giving us

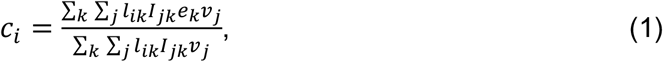

where 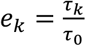 is the enrichment of heritability for topic *k* annotation (*A*_*k*_). *e*_*k*_ is estimated from S-LDSC for each annotation *A*_*k*_ (one topic at a time), where the estimation is denoted as *ê*_*k*_ with standard error *w*_*k*_. Assuming *l*_*ik*_ is known, we use the estimated cell-level enrichment 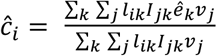 as the cell level score. The variance of *c*_*i*_ is given by 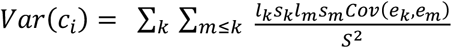 where *s*_*k*_ = ∑_*j*_ *I*_*jk*_ and *S* = ∑_*k*_ *l*_*k*_ *s*_*k*_. S-LDSC regresses the variants’ GWAS summary statistics against their LD scores with respect to *A*_*k*_ to get *ê*_*k*_ and standard error *w*_*k*_. The LD score of a variant *p* with respect to annotation *A*_*k*_ is given by 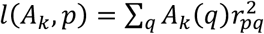, where *A*_*k*_(*q*) is the annotation *k* for variant *q* and *r*_*pq*_ is the correlation between variants *p* and *q*. Thus, we have the covariance *Cov*(*e*_*k*_, *e*_*m*_) = *w*_*k*_*w*_*m*_*Cor*(*l*(*A*_*k*_), *l*(*A*_*m*_)), where *Cor*(*l*(*A*_*k*_), *l*(*A*_*m*_)) is the correlation between LD scores with respect to annotation *A*_*k*_ and *A*_*m*_ across variants. To simplify implementation, we approximated *Cor*(*l*(*A*_*k*_), *l*(*A*_*m*_)) using *Cor*(*A*_*k*_, *A*_*m*_) when calculating the cell score variance.

With the cell score and variance, SCADS calculates a *z* score and *p* value for each cell. The BH procedure is used for controlling FDR. A limitation of the current cell score variance calculation is that we do not account for the uncertainty of *l*_*ik*_ and *I*_*jk*_, therefore the actual cell score variance should be larger. In practice, we found that the cell score variance is mainly driven by the S-LDSC estimate error *w*_*k*_ and the *p*-values are well calibrated based on simulations studies. To further stabilize this score and prevent bias from topics with large estimation errors, we apply a shrinkage method (ASH^66^) on S-LDSC estimates (*ê*_*k*_ and *w*_*k*_) with a uniform prior to obtain posterior mean estimates of *ê*_*k*_ for SCADS cell score *ĉ*_*i*_ calculation. As it has been shown for small annotations, S-LDSC tends to have inaccurate error estimation^67^, thus SCADS excludes any annotation in the cell score calculation if it is < 0.5% of the genome.

### Simulation of scATAC-seq Data I

For Fig. 2 and Supplementary figure 1 related simulations, we simulated scATAC-seq data based on a Poisson NMF model,

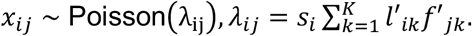

The Poisson rate (*λ*_*ij*_) in cell *i* and region *j* is proportional to *s*_*i*_, the total read count in cell *i*, and is a mixture of *K* rates. The *k*th rate in region *j, f*′_*jk*_, represents the Poisson rate of observing a read in region *j* for the *k*th cell type. *l*′_*ik*_ represents the mixture weights; in our simulation, we set 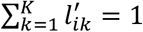. This model can be readily transformed into a topic model: *l*′_*ik*_ represents the grade of membership for cell *i* (same as *l*_*ik*_) and *f*′_*jk*_ normalized by the sum across all regions in topic *k* will give the probability of observing a read in topic *k* for region *j* (*f*_*jk*_).

We set ***f***′_*k*_ = (*f*′_1*k*_, …, *f*′_*jk*_) matching the chromatin accessibility of a real cell type. Then we designated *K* = 5 and performed peak calling on the pseudobulk data for 5 cell types (Monocytes, CD4^+^ T cell, CD8^+^ T cells, NK cells, B cells) from a scATAC-seq dataset^30^. We randomly selected 260K regions that are peaks in at least one cell type (*J* = 260,000). For a closed region *j* in cell type *k*, we drew *f*′_*jk*_ from a log normal distribution, i.e. log 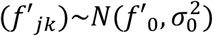; for an open region we used log 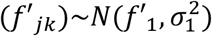. We assigned the ratio of *f*′_1_ to *f*′_0_ as 7.25, matching the ratio of read depth in peak and non-peak regions in the pseudobulk data. Additionally, we set 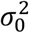 and 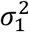 to achieve varying coefficients of variance (CV) of *f*′_*jk*_. Specifically, for 0% CV, we would set 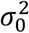 and 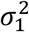 to 0, and the Poisson rate for open and close regions in a topic are constants.

If we restricted *l*^′^_*ik*_ to either 0 or 1, then we would be assuming there are only *K* distinct cell states. To allow for continuous cellular state transitions in our simulation, we drew loadings for cell *i*, ***l***′_*i*_ = (*l*′_*i*1_, …, *l*′_*i*-_)^⊤^, independently from a Dirichlet distribution, 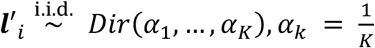, for all *k* = 1, …, *K*. We had set *f*′_0_ so that 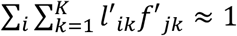, so *s*_*i*_ controls the read depth per cell. We simulated *s*_*i*_ from a log-normal distribution (mean = 20,000, standard error = 0.173), matching cell read depth for typical scATAC-seq studies^18^.

### Simulation of GWAS Data

We simulated quantitative traits using individual level genotype data and calculated GWAS summary statistics. First, we used UKBiobank genotype data comprising approximately 6.23 million variants for 45,087 individuals. The genotypes were processed as previously described^68^. We generated *K* variant level annotations for the *K* cell types used in simulating scATAC-seq data above. The *k*th annotation includes all open regions in cell type *k*. We simulate the quantitative trait *y* using an additive model *y* = *Σ*_*p*_*X*_*p*_*β*_*p*_^0^ + *Σ*_*p*_*Σ*_*k*_*X*_*p*_*β*_*pk*_ + *ε*. Here, *X*_*p*_ is the genotype of variant *p. ε* ∼ *N*(0, *σ*^2^) is assumed to be independent across individuals. *β*_*p*_^0^ is the effect size of variant *p*, and *β*_*pk*_ is the effect size depending on the *k*th cell type derived annotation. We simulate *β*_*p*_^0^and *β*_*pk*_s independently following mixture distributions, *β*^0^_*p*_ ∼ *π* ^*b*^*N*(0, *σ*^2^) + (1 − *π* ^*b*^)*δ*_0_ and *β*_*pk*_ ∼ *π*_*k*_ *N*(0, *σ*^2^) + (1 − *π*_*k*_)*δ*_0_. *δ*_0_ is the point mass at 0. *π*^*b*^ is the baseline fraction of causal variant, and it does not depend on annotations. *π*_*k*_ is the additional fraction of causal variants in annotation *k*. If the annotation is derived from a causal cell type, then we set *π*_*k*_ >0. In our simulations, we have set it as 10x or 20x of *π* ^*b*^, matching the estimated enrichment from real data. If the cell type is non-causal, we assigned *π*_*k*_ =0. *σ* is the average causal effect and is determined based on heritability. We experimented with a few different values for *π* ^*b*^ (e.g. 1.5 × 10^−4^, 2.5 × 10^−4^ and 1 × 10^−3^) and for heritability (e.g. 0.25 or 0.5) to represent different genetic architecture of complex traits.

### Null Simulation

We simulated scATAC-seq data for 1200 cells and 130,000 peaks from three cell types (Monocytes, CD4^+^ T cells, CD8^+^ T cells) as described above. We simulated GWAS summary statistics independently for 200 traits, with *π*_*b*_ = 2.5 × 10^−4^ and *π*_*k*_ = 0, *k* = 1, 2, 3. The total heritability for a trait is set to 0.5.

### Simulation of scATAC-seq Data II

For Fig. 3 related simulations, we implemented an alternative simulation procedure as described in the SCAVENGE paper^26^. We simulated 1,000 cells (500 monocytes and 500 B cells) with 30,000 reads per cell (*s*) and a 30% background noise rate (*r*). Other inputs included: (1) a pseudobulk accessibility matrix providing read counts per genomic peak for each cell type using a reference (See Data for simulating cell types in Data Availability section) and (2) a corresponding set of genomic peak coordinates. Each simulated cell was uniquely labeled according to its cell type and index (e.g., *Mono_1, Bcell_2*). Following the SCAVENGE paper, for each cell type, the probability of accessibility at each peak was estimated by normalizing the bulk read counts across peaks and scaling these probabilities according to the desired sequencing depth ***s*** and noise rate ***r***. For each cell type *c*, the accessibility probability of peak *j* was computed as:

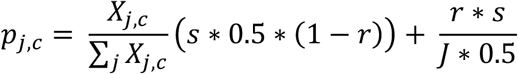

where *X*_*j*,c_ is the read count for peak *j* in cell type *c* and *J* is the number of peaks. To avoid extreme sampling behavior, probabilities were capped at a maximum value of 0.9. We then simulated accessibility for each peak in each cell by performing two independent Bernoulli draws to account for both haplotypes. The resulting values were summed to yield accessibility counts of 0, 1, or 2 per peak.

### SCADS running parameters

SCADS uses the baseline LD model (version 2.2, hg19) to run S-LDSC. For GWAS and scATAC-seq data in genome build other than hg19, we first lift them over to hg19 via LiftOver^69^.

For the *K* selection when running SCADS, we generally choose a number that roughly matches the number of cell types or clusters. In simulations, we found SCADS results are robust to the choice of *K* topics. In simulations, we have tried to run SCADS with *K*=3, 6, 9 when there are 5 true underlying topics. We found the results are consistent among runs with different *K*s and enriched cells were recovered by SCADS though specifying more topics is preferred (Supplementary Fig. 6). We specified *K* = 25 for the hematopoiesis data in **Fig. 5** and *K* = 15 for colon tissue data in **Fig. 6**.

### Running of comparator software

We compared SCADS with g-chromVAR and SCAVENGE, both of which required a list of fine-mapped variants as input. For the monocyte count trait, we utilized a list of fine-mapped variants provided by SCAVENGE. And for all other traits, we performed Sum of Single Effects (SuSiE)^70^ fine-mapping (L=1) to determine the list of causal variants (PIP>0.001). To run g-chromVAR, we first corrected the single-cell chromatin accessibility matrix for GC bias and constructed a matched background peak set to control for technical variation. We then computed weighted deviation *z*-scores for each cell using genome-wide trait enrichment weights (i.e. fine-mapped posterior probabilities). SCAVENGE utilized these z-scores to identify seed cells and constructed a cell-cell similarity network using a mutual k-nearest neighbor (M-KNN) graph with k =30. Network propagation was then executed using a random walk with restart (RWR) algorithm (gamma = 0.05) starting from the defined seed cells to calculate network propagation score. Outliers in the NP scores were capped at the 95th percentile and min-max scaled to generate the final trait relevance score.

### Motif analysis by HOMER

We performed motif enrichment using HOMER (v5.1)^71^. Motif discovery was performed using *findMotifsGenome*.*pl* with the ‘–size given’ argument to use the exact peak widths from the input BED files.

### Differential accessibility analysis within CD8^+^ CM T cells

We grouped CD8^+^ CM T cells into SCADS higher-scoring group (score > median of all CD8^+^ CM T cells) and SCADS lower-scoring group (score < median of all CD8^+^ CM T cells). We performed differential accessibility between these two groups for all regions with > 5 reads aggregated across all CD8^+^ CM T cells. We ran DESseq2 to call differential accessible regions with dispersion parameter of 0.01 and FDR < 0.05.

### Bayesian fine-mapping using topic annotations

We used topic annotations with significant enrichment (*p* < 0.05) from SCADS to perform functional annotation informed fine-mapping via PolyFun^72^ and SuSiE. SuSiE was run with L=5 for all LD blocks as previously defined^73^.

### Correlation between variant accessibility and SCADS score

After obtaining SCADS scores, the cell population of interest (e.g., CD8^+^ CM T cells) was selected and categorized into five quantile-based score bins. For each fine-mapped variant, we counted the reads that fell into the 500bp region around the variant for all cells in a bin as the bin accessibility. We then converted the bin accessibility to counts-per-million (CPM), defined as 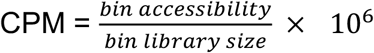, where the bin library size is all reads in cells mapped to the bin. We retained variants with CPM≥ 5 in at least two bins and calculated the Pearson correlation between CPM and SCADS scores across bins for each variant.

### Regulon enrichment

We obtained regulons for cell types in the hematopoiesis system as described before^40^. In brief, SCENIC^39^ was employed on a scRNA-seq data of 2.15M peripheral blood mononuclear cells to identify 376 TF regulons and their respective activities for 7 major cell types (B cells, CD4^+^ T cells, CD8^+^ T cells, NK cells, dendritic cells, monocytes, and other T cells). The SCENIC framework, which integrates co-expression analysis with TF binding motif enrichment in cis-regulatory regions of candidate target genes, ensures that the target genes within a regulon harbor a predicted binding site for the cognate TF.

We retained fine-mapped variants with an absolute CPM and SCADS score correlation of greater than 0.2. For each variant, we retrieved the nearest protein-coding gene (or genes, in the case of ties) based on genomic distance from the hg19 transcript reference (org.Hs.eg.db), restricting to standard autosomes, removing noncanonical or unplaced contigs. The resulting gene list contained the target gene set for downstream enrichment testing. For each TF regulon, we constructed a 2×2 contingency table containing the number of (a) target genes in the regulon, (b) non-target genes in the regulon, (c) total genes in regulon, and (d) background genes not in regulon. A one-sided Fisher’s exact test (alternative = “greater”) was performed to determine whether the overlap between the target gene set and the TF regulon was greater than expected by chance. The odds ratio from the test is reported as the enrichment. Multiple testing correction was applied using the Benjamini–Hochberg (BH) procedure to control the false discovery rate (FDR).

### Culture of Colon Organoids

We chose to utilize colon organoids as the validation model for our findings derived from intestinal epithelial cells, based on their availability and the similarity of their epithelial cell types and functional characteristics to the small intestine. The human colon organoid line used for CRISPRi experiments was derived from a de-identified sigmoid colon tissue specimen obtained with informed consent under IRB-STUDY02000180 at Dartmouth Health and IRB-STUDY00033600 at Dartmouth College. The sample was inspected by the surgeon and pathologist assistant and were later confirmed to be histologically normal by H&E staining.

Normal colon tissue was washed in cold PBS. The epithelium was stripped from the muscularis and minced into small pieces with a scalpel. Tissue fragments were incubated in 2mg/mL collagenase/dispase for 20 minutes at 37 °C, mixed 10 times with a 10 mL serological pipette and incubated for another 20 minutes, again at 37 °C, before further triturating and diluting with 10 mL of ice cold Advanced DMEM F12. The crypt containing supernatant was then transferred to a new 15mL tube and centrifuged at 80×g for three minutes and the supernatant carefully aspirated. The loose crypt pellet was gently washed a second time with Advanced DMEM F12 and inspected under the microscope. After a final centrifugation at 200×g for 2 minutes, crypts were embedded in Matrigel basement membrane extract and a total of 50 µl was plated in three drops per well of a 24 well plate and placed in the incubator for 10 minutes for polymerization. Half a milliliter of pre-warmed expansion media was added per well.

Organoids were maintained in an expansion media formulation tailored to human colon organoids as described^74^ with some minor modifications. A basal medium comprised of Advanced DMEM F12, Glutamax (1×), HEPES (15mM), and 100 U/ml penicillin-streptomycin (referred to as AdvDMEM+++) was further supplemented with B-27 Plus Supplement (1×), 1.25 mM *N*-acetylcysteine, 0.5 nM NGS-Wnt-Fc, 20% RSPO1 conditioned media, 100ng/mL rhNoggin, 0.5 µM A83-01, 50 ng/mL EGF, 25 nM Leugastrin, 50 ng/mL FGF-2, 100 ng/mL IGF-1. To passage for general maintenance and expansion, Day 7 hCOs were extracted from Matrigel using ice cold AdvDMEM+++ and dissociated with 0.5x TryplE Select for 3 minutes at 37°C. Digested hCO fragments were replated in Matrigel at a split ratio of 1:6 to 1:10. Expansion media was supplemented with 50 nM Chroman-1 for the first 2 days after passage. To differentiate hCOs, cultures were digested to single cells and plated at low density in the above media formulation with the addition of 1 ng/mL NRG1 for 4 days before changing to Balance media (Day 4 to 11) which includes the following substitutions to aid in mature cell type specification: 0.1 nM NGS-Wnt-Fc, 10% RSPO1 conditioned media, 5 ng/mL EGF, 10 ng/mL NRG1, and the removal of A83-01. Balance media was replaced every 2-3 days.

### CRISPRi of rs7553638

The lentiCRISPRi(v2)-Blast plasmid was a gift from Neville Sanjana (Addgene plasmid # 170068)^75^. Briefly, 2nd generation lentivirus was produced by polyethylenimine transfection of HEK293T cells with psPAX2 and pMD2.G (a gift from Didier Trono). Viral supernatants were harvested, clarified by centrifugation, passed through a 0.45nm filter, and concentrated by ultracentrifugation at 70,000 x g for 2 hours at 17C as previously described^76^. The viral pellet was resuspended in colon organoid expansion media. An organoid line stably expressing KRAB-dCas9-MeCP2 was generated by lentiviral infection at low multiplicity of infection, followed by continuous selection with 5 ug/mL blasticidin. Seven days after transduction, outgrowing organoids were digested to single cells with TryplE, filtered with a 20um filter and plated at low density in matrigel. Isogenic organoids were manually picked under a microscope and split into 2 wells of a 48w plate. Organoids from one of the clonal wells were harvested for immunofluorescence screening of HA-tagged KRAB-dCas9-MeCP2 expression. Two clones were identified to have specific and ubiquitous expression of the HA tagged fusion and were expanded and cryopreserved for downstream assays. To confirm the functionality of the CRISPRi organoid lines, KRAB-dCas9-MeCP2 isogenic organoids were subsequently transduced with lentivirus encoding an sgRNA targeting the promoter region of either CD46 or B2M and conferring puromycin resistance. 48 hours later, organoids were selected with 2ug/mL puromycin and harvested at day 10 post infection for immunofluorescent staining of cell surface CD46 or B2M to confirm knockdown efficiency.

### rs7553638 sgRNA design and cloning

Two sgRNA targeting within 20bp of the rs7553638 variant were selected based on proximity and for the best possible off-target^77^ and on-target score^78^. The sequence for rs7553638-sg1 is *GCATCATCAGCACCTGTGCC* and rs7553638-sg2 is *GACGATGACTGCAAATACTG*. Guides were cloned as previously described into sgRNA expression vector pJR101^79^. The pJR101 plasmid was a gift from Marco Jost & Jonathan Weissman (Addgene plasmid # 187241). Briefly, sgRNA sequences were appended with flanking sequences containing BstX1/BlpI overhangs and two complimentary oligonucleotides were synthesized (IDT) phosphorylated and annealed and ligated into BstX1/BlpI digested pJR101 according to standard molecular cloning techniques.

### Single-cell RNA sequencing of colon organoid and cell typing

The plasmids encoding rs7553638-sg1 and rs7553638-sg2 were pooled with another Perturb-seq library for cost sharing purposes. A total of four non-targeting control sgRNA from the GeCKOv2 library(PMID: **25075903)** were included as negative controls. Non-targeting sgRNA protospacer sequences were as follows: sgNTC-1 *CGCGGAAATTTTACCGACGA*, sgNTC-2 *CGTGTGTGGGTAAACGGAAA*, sgNTC-3 *GCCTGCCCTAAACCCCGGAA*, sgNTC-8 *GCACGCTGTACAGACGACAA*. Plasmids were pooled at equimolar ratio and lentivirus was generated as described above. A functional titer was performed by serial dilution on HEK293T cells and assessed by the GFP reporter expression. For library transduction, dCAS9 expressing organoids were dissociated into single cells by incubation in 1x TryplE Select supplemented with Chroman-1,for 15 minutes with pipetting every 5 minutes. Dissociated cells were passed through a 30-micron filter (Pluriselect) to remove clumps and debris. To achieve an MOI = 5, 250k cells were spinoculated for 1hr at 600xg and 30C with 50ul of concentrated lentiviral library in a total volume of 300ul expansion medium in 2 wells of a 24w plate supplemented with Chroman-1 and 6ug/mL polybrene. After spinoculation, the plate was transferred to a 37C incubator for an additional 5 hours. Subsequently, cells were collected and plated into Matrigel and overlaid in Expansion Media supplemented with 1 ng/mL NRG1 for 4 days followed by a switch to balance medium for days 4-11 to induce differentiation. Two days after transduction, 2ug/mL of puromycin was added to select for cells expressing sgRNA. Differentiated colon organoids were collected on Day11 in ice cold AdvDMEM+++ pelleted and and dissociated to single cells by resuspension in 2mL prewarmed 0.05% Trypsin + EDTA for ∼10min with frequent pipetting, then washed by dilution with 12mL of AdvDMEM+++. Cells were resuspended in 0.04% BSA in PBS and clumps were removed by passing through a 30-micron filter. Cells were pelleted and passed through a subsequent 20-micron filter to further isolate single cells. Single cells were counted and viability assessed by LUNA FX7. Five hundred thousand live cells were submitted to the Genomics Shared Resource at Dartmouth Cancer Center for scRNA-sequencing with 10X Genomics Chromium Single Cell Flex Gene Expression Assay. Flex Gene Expression was performed according to 10X genomics protocols (CG000782,CG000787) and custom CRISPR probes were designed (technical note CG000814). Cells were fixed then split equally among a 4-plex hybridization. A custom CRISPR probe library complementary to the guide protospacer sequences was designed and spiked into the hybridization step. Fifteen thousand cells per plex were mixed and loaded into a single lane for emulsion with the ChromiumX partitioner. CRISPR libraries and transcriptomic libraries were prepared separately according to standard GEM-X Flex protocols before sequencing on the Illumina NextSeq2000. Gene expression libraries were sequenced to ∼27,000 mean reads per cell which achieved ∼4,600 median genes per cell. The CRISPR library was sequenced to ∼9,000 mean reads per cell.

### Differential gene expression analysis for CRISPRi

We analyzed the CRISPRi screen data with the SCEPTRE^80^ R package, supplying the gene-expression matrix, the binary gRNA matrix, a guide-target annotation table, and specifying MOI as “high.” Cell clusters generated via Seurat were annotated using known markers^81^ and included as additional covariates to default covariates including library size (log-transformed RNA and gRNA UMIs), molecular complexity (log-transformed non-zero RNA and gRNA features), and mitochondrial content. Following the SCEPTRE workflow, we first created a SCEPTRE object, defined the analysis parameters, and assigned guides to individual cells via a mixture model. Quality-control, calibration, and statistical power diagnostics were assessed to ensure reliable inference. After which we performed the discovery analysis to test for differential expression of each gRNA’s target gene, yielding calibrated p-values and effect size estimates while properly accounting for multiplexed perturbations.

## Data availability

### Data for simulating cell types

scATAC-seq read count matrix summary for human immune cells is obtained from Gene Expression Omnibus (GEO, https://www.ncbi.nlm.nih.gov/geo/) with ID (GSE74912).

### Genotypes data for simulating phenotype

From UK Biobank as described previously^68^.

### scATAC-seq data

All single-cell datasets used in the paper are publicly available. Hematopoiesis scATAC-seq datasets were downloaded from https://github.com/GreenleafLab/MPAL-Single-Cell-2019. Colon tissue^54^ scATAC-seq datasets downloaded from https://doi.org/10.5061/dryad.0zpc8672f.

### GWAS summary statistics

GWAS traits of Monocyte count (https://www.ebi.ac.uk/gwas/studies/GCST90002393), IBD (https://www.ebi.ac.uk/gwas/studies/GCST004131), Asthma (https://www.ebi.ac.uk/gwas/studies/GCST010042), Type-1 diabetes (https://www.ebi.ac.uk/gwas/studies/GCST90014023), Hypothyroidism (https://www.ebi.ac.uk/gwas/studies/GCST90319320), Rheumatoid Arthritis (https://www.ebi.ac.uk/gwas/studies/GCST90018910), Primary biliary cholangitis (https://www.ebi.ac.uk/gwas/studies/GCST90061440) were downloaded from GWAS catalog^82^. The Lupus(https://www.nature.com/articles/s41467-023-36306-5) and Eczema/Atopic dermatitis (https://www.nature.com/articles/s41467-025-58310-7) were downloaded from their respective sources.

## Code availability

R package and installation instructions are provided on GitHub (https://github.com/szhaolab/scads). Accompanying analysis scripts are also deposited on GitHub (https://github.com/szhaolab/scads_paper)

## Acknowledgement

This work was supported by P20GM130454 and R35GM154925 (to S.Z.). The Munck-Pfefferkorn Education and Research Fund at Dartmouth Cancer Center. A T32 training grant (5T32CA260262-07) to L.Y.. We like to acknowledge Dr. Xin He at University of Chicago for discussion of the manuscript; Dr. Lucas Salas at Dartmouth College and Dr. Hongbo Liu at University of Rochester for discussions regarding scATAC-seq data.

