## Supplemental Materials for "Connecting polygenic disease risk to cell states and regulatory programs through single-cell chromatin accessibility"

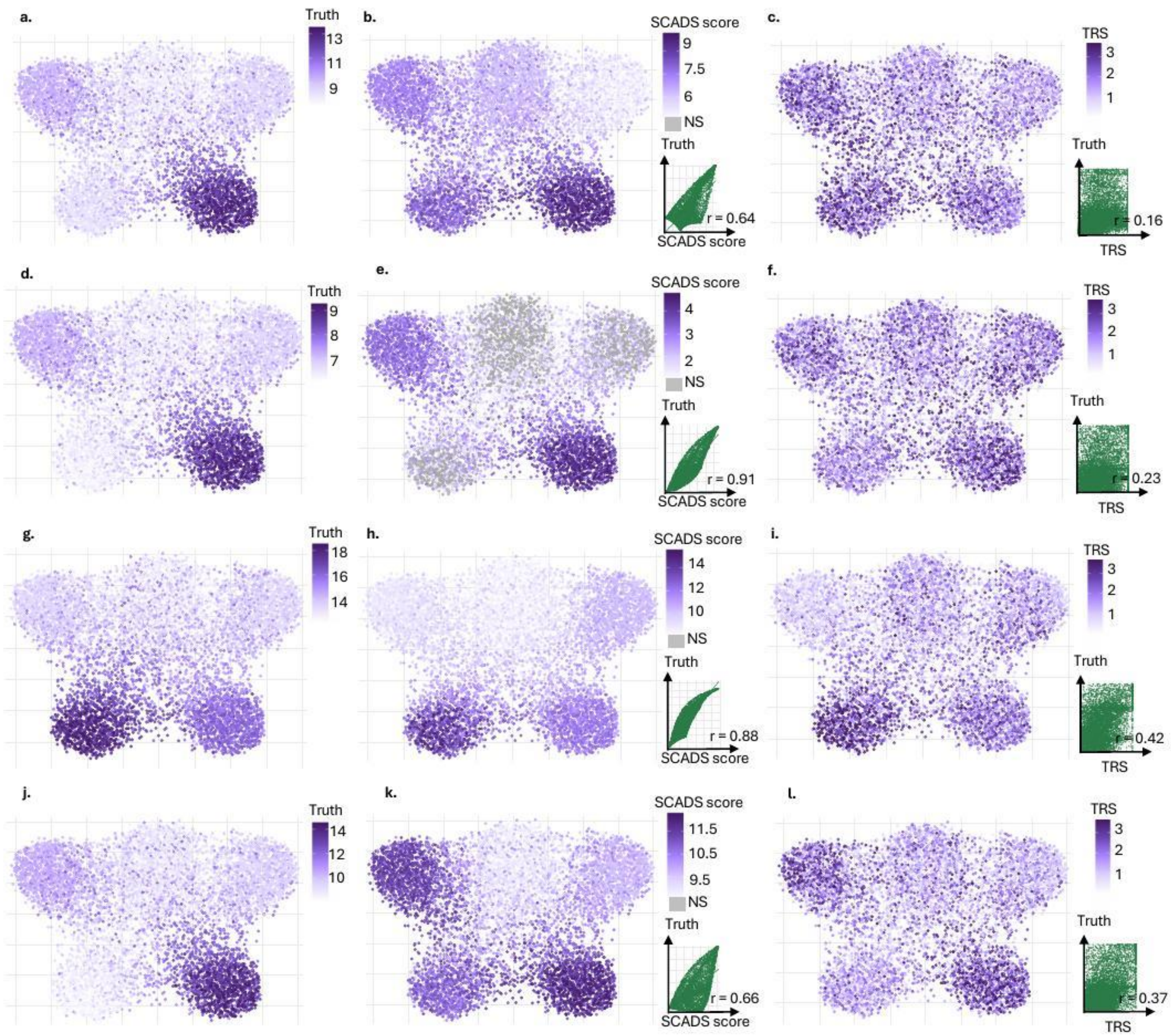


**Supplementary Figure 1. Comparative performance of SCADS and SCAVENGE across additional simulation settings.** These plots follow the format of **Fig. 2.** The simulated scATAC-seq data comprises 260K peaks across 7.5K cells, contains five topics and an average of 20K reads per cell. The ratio of reads in open versus closed regions is set to 7.25 with a CV of 20% (see Methods). **a–c**. Results for simulated GWAS with $\pi^b = 2.5 \times 10^{-4}$, one causal topic (fraction of causal variant in the casual topic open regions is 20x of $\pi_b$ ), and a total trait heritability of 0.25. **d-f.** Results for simulated GWAS with $\pi^b = 2.5 \times 10^{-4}$, one causal topic (fraction of causal variant in the casual topic open regions is 10x of $\pi_b$ ), and a total trait heritability of 0.5. **g–i.** Results for simulated GWAS with $\pi^b = 2.5 \times 10^{-4}$, two causal topics (fraction of causal variant in the casual topic open regions is 20x of $\pi_b$, each), and a total trait heritability set to 0.5. **j-l.** Results for simulated GWAS with $\pi^b = 1.5 \times 10^{-4}$, one causal topics (fraction of causal variant in the casual topic open regions is 20x of $\pi_b$), and a total trait heritability of 0.5. SCADS scores with FDR > 0.05 (non-significant or NS) are colored in grey.

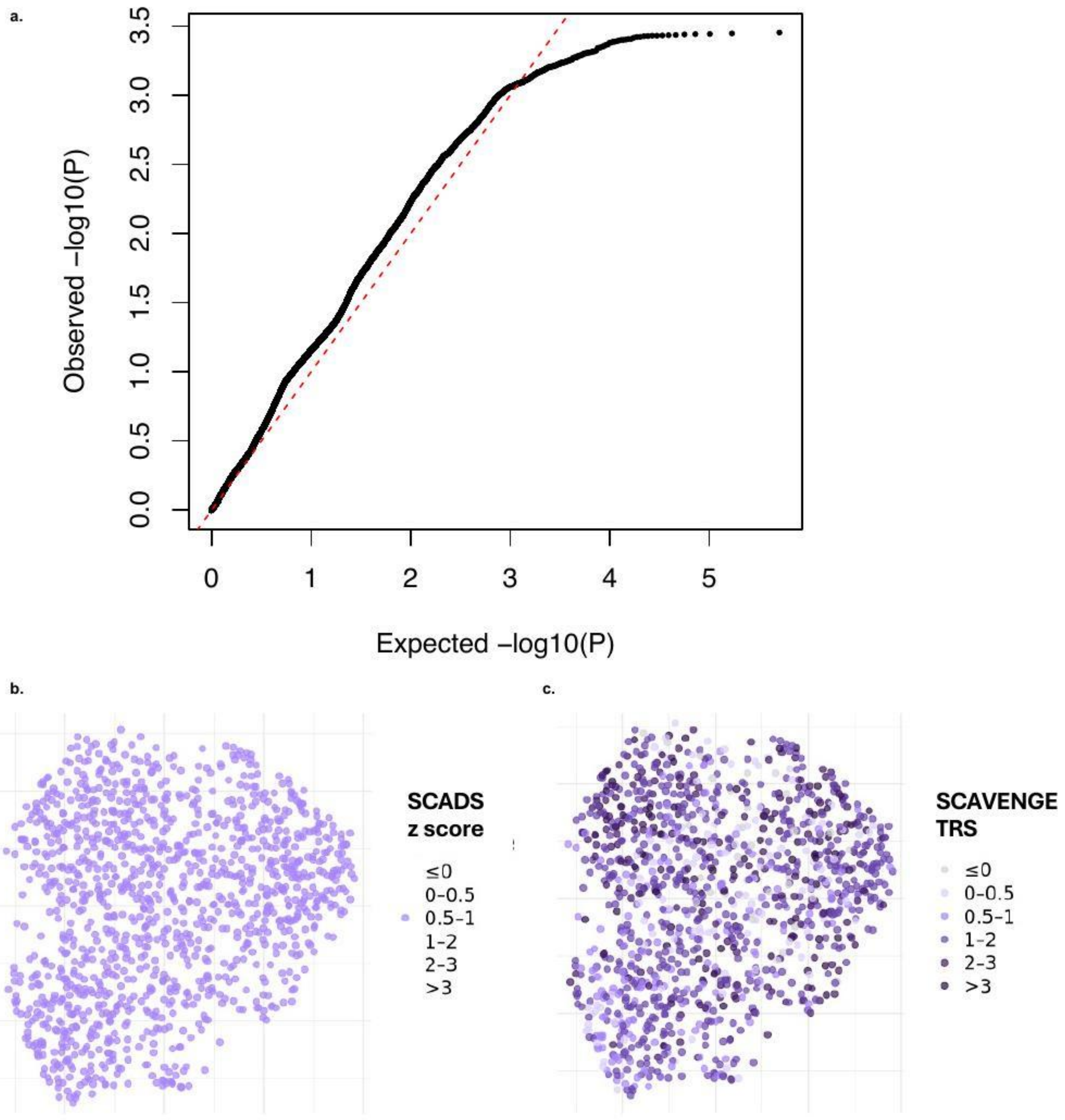


**Supplementary Figure 2. Evaluation of SCADS and SCAVENGE in null simulation. a**, Quantile-quantile (Q-Q) plot displaying SCADS *p*-values (y-axis) against a uniform distribution (x-axis) for the null simulation. **b–c**, UMAP visualizations of a representative simulated GWAS trait, colored based on **b**, SCADS z-scores (all BH adjusted p-values > 0.05) and **c,** SCAVENGE scores.

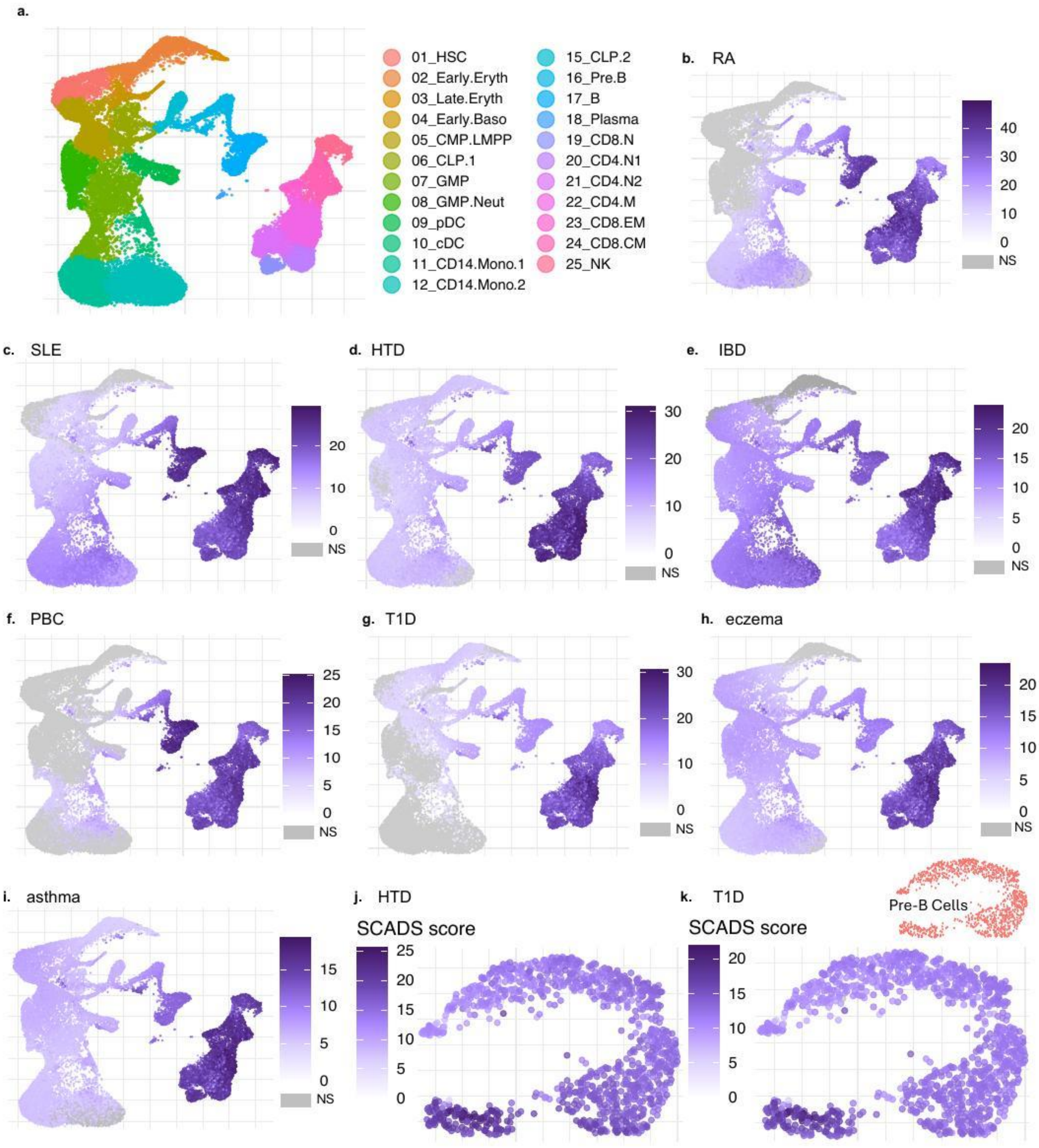


**Supplementary Figure 3. SCADS results on human hematopoiesis cells for eight autoimmune traits. a,** UMAP visualization of scATAC-seq data from the human hematopoietic system, with clusters labeled according to cell type annotations from the reference study[1]. **b–i,** SCADS scores for eight autoimmune conditions: **b,** rheumatoid arthritis (RA); **c**, systemic lupus erythematosus (SLE); **d,** hypothyroidism (HTD); **e,** inflammatory bowel disease (IBD); **f,** primary biliary cholangitis (PBC); **g,** type 1 diabetes (T1D); **h,** atopic dermatitis (eczema), and **i,** asthma. Cells with non-significant enrichment (FDR > 0.05) are colored in grey. **j-k**, SCADS scores for **j,** HTD and **k,** T1D for pre-B cells only.

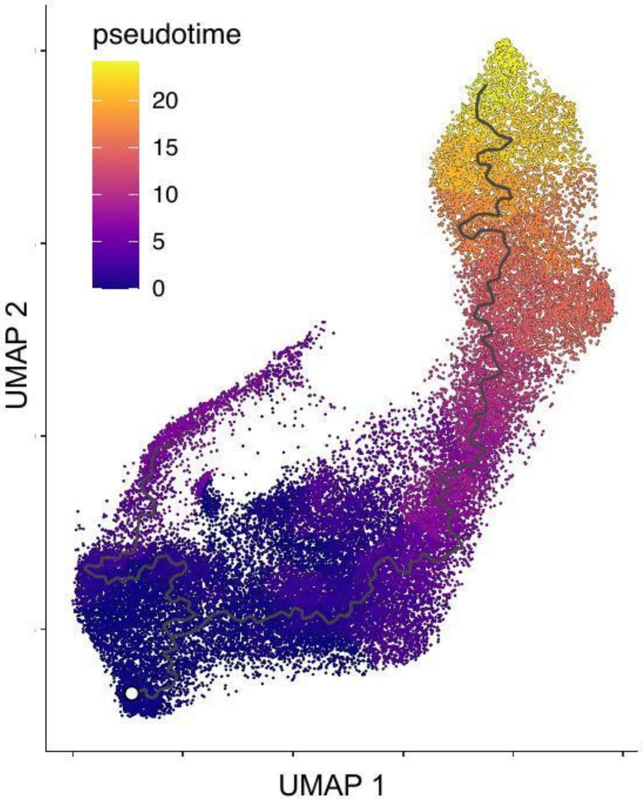


**Supplementary Figure 4. Pseudotime trajectory of the enterocyte differentiation lineage.** UMAP visualization of the colon epithelial cells colored by pseudotime value, highlighting the differentiation progression from intestinal stem cells to mature enterocytes. The reconstructed trajectory originates at the stem cell population (white circle) and extends along two branches, with the longer primary branch (right) representing the full differentiation path. Pseudotime was assessed using Monocle3[2,3], in which a principal graph was first inferred within the UMAP embedding and then refined to a linear backbone by extracting the shortest paths between manually defined anchor nodes while pruning auxiliary branches using the igraph[4] (v2.1.4) R package.

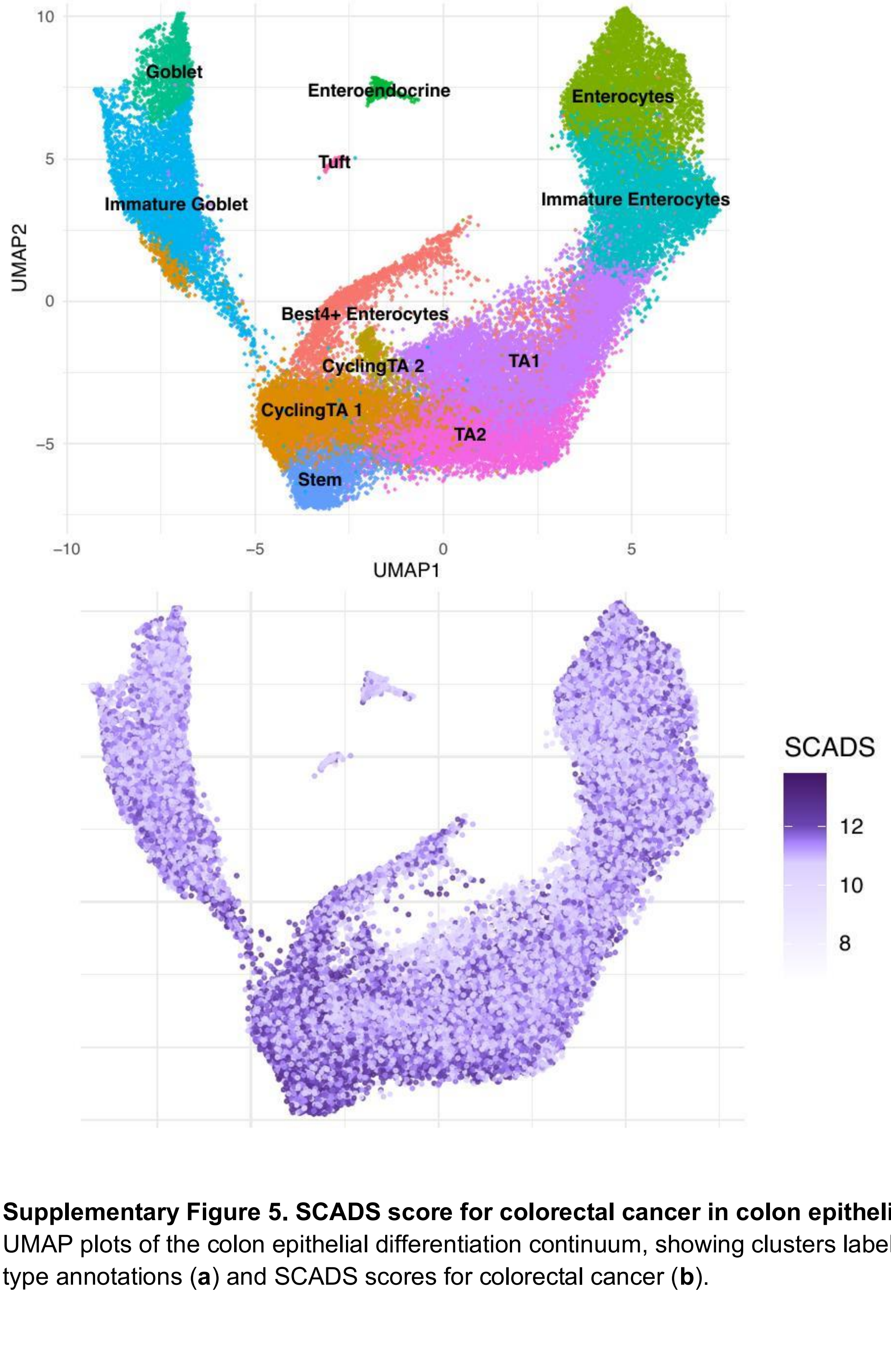


**Supplementary Figure 5. SCADS score for colorectal cancer in colon epithelial cells. a–b.** UMAP plots of the colon epithelial differentiation continuum, showing clusters labeled by cell type annotations (**a**) and SCADS scores for colorectal cancer (**b**).

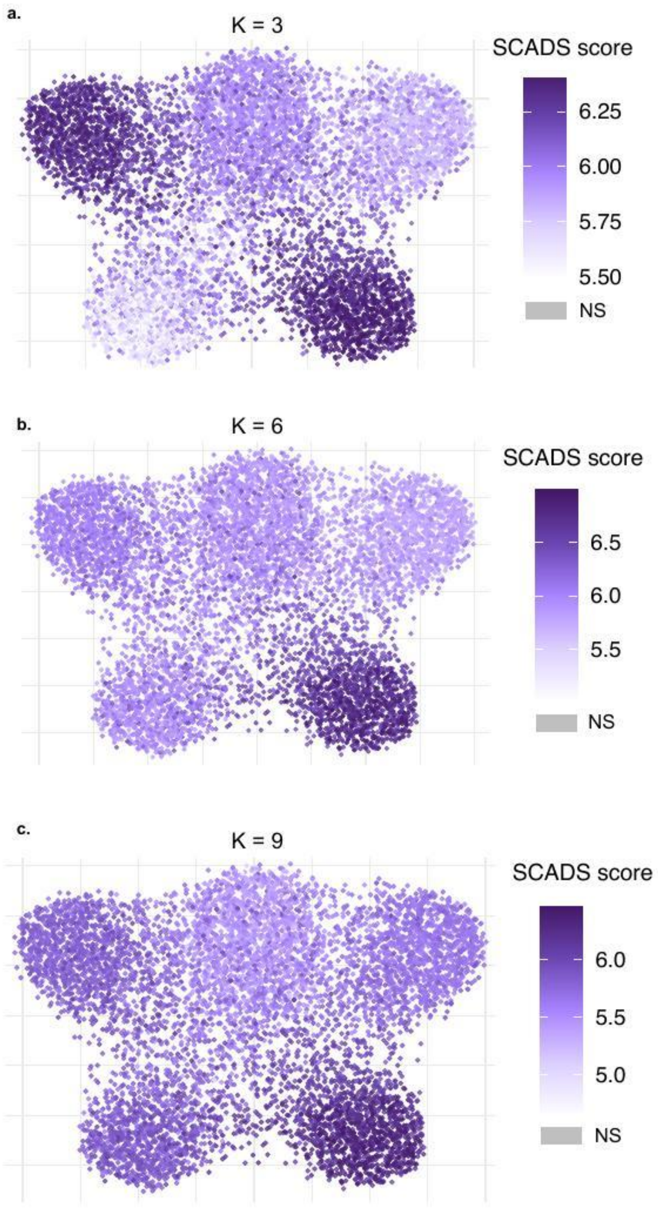


**Supplementary Figure 6. Robustness of SCADS to the choice of number of topics.** These UMAP plots present results from the dense setting described in **Fig. 2**. We evaluated the impact of varying the number of topics (K): **a,** K = 3, **b,** K = 6, and **c,** K = 9.

**Supplementary Table 1. Motif enrichment for topic 8 by HOMER.**
A total of 301 transcription-factor binding motifs passed the significance threshold (q < 0.05) in HOMER. The table here listed those with top enrichment (ratio between % target sequence with motif and % background sequence with motif > 1.5). The table is sorted in ascending order of *p*-values. All motifs in this table have *q* value < 0.0001.

| Motif Name | Consensus | log p-value | # of Target Sequences with Motif(of 100297) | % of Target Sequences with Motif | # of Background Sequences with Motif(of 99045) | % of Background Sequences with Motif | Enrichment |
|---|---|---|---|---|---|---|---|
| CTCF | AYAGTGCCMYCTRGTGGCCA | -1.25E+04 | 12818 | 12.78% | 2198.8 | 2.22% | 5.76 |
| BORIS | CNNBRGCGCCCCCTGSTGGC | -7.19E+03 | 14922 | 14.88% | 4826.2 | 4.87% | 3.06 |
| ETS1 | ACAGGAAGTG | -4.31E+03 | 34062 | 33.97% | 21082.9 | 21.27% | 1.60 |
| Fli1 | NRYTTCCGGH | -4.29E+03 | 36431 | 36.34% | 23121.5 | 23.32% | 1.56 |
| Etv2 | NNAYTTCCTGHN | -3.90E+03 | 31052 | 30.97% | 19103.5 | 19.27% | 1.61 |
| ETV4 | ACCGGAAGTG | -3.75E+03 | 35761 | 35.67% | 23305.4 | 23.51% | 1.52 |
| GABPA | RACCGGAAGT | -3.47E+03 | 29415 | 29.34% | 18326.7 | 18.49% | 1.59 |
| Ets1-distal | MACAGGAAGT | -3.19E+03 | 11841 | 11.81% | 5256.9 | 5.30% | 2.23 |
| EWS:ERG-fusion | ATTTCCTGTN | -2.61E+03 | 23651 | 23.59% | 14784.7 | 14.91% | 1.58 |
| EWS:FLI1-fusion | VACAGGAAAT | -2.51E+03 | 19486 | 19.44% | 11569 | 11.67% | 1.67 |
| Elk1 | HACTTCCGGY | -2.35E+03 | 19827 | 19.78% | 12064.1 | 12.17% | 1.63 |
| Elk4 | NRYTTCCGGY | -2.23E+03 | 19674 | 19.62% | 12098.6 | 12.20% | 1.61 |

| | | | | | | | |
|---|---|---|---|---|---|---|---|
| IRF2 | GAAASYGAAASY | -2.10E+03 | 5987 | 5.97% | 2285.4 | 2.31% | 2.58 |
| PU.1 | AGAGGAAGTG | -2.04E+03 | 16282 | 16.24% | 9671.8 | 9.76% | 1.66 |
| RUNX | SAAACCACAG | -1.94E+03 | 21006 | 20.95% | 13632.4 | 13.75% | 1.52 |
| IRF1 | GAAAGTGAAAGT | -1.87E+03 | 7064 | 7.05% | 3118.9 | 3.15% | 2.24 |
| ELF1 | AVCCGGAAGT | -1.86E+03 | 17991 | 17.94% | 11297.4 | 11.40% | 1.57 |
| ETS Promoter | AACCGGAAGT | -1.76E+03 | 12350 | 12.32% | 6993.3 | 7.05% | 1.75 |
| ISRE | AGTTTCASTTTC | -1.66E+03 | 4036 | 4.03% | 1395.7 | 1.41% | 2.86 |
| ETS:RUNX | RCAGGATGTGGT | -1.61E+03 | 4840 | 4.83% | 1894.6 | 1.91% | 2.53 |
| IRF8 | GRAASTGAAAST | -1.61E+03 | 12390 | 12.36% | 7224.1 | 7.29% | 1.70 |
| IRF3 | AGTTTCAKTTTC | -1.47E+03 | 12561 | 12.53% | 7539.4 | 7.61% | 1.65 |
| Fosl2 | NATGASTCABNN | -1.25E+03 | 9059 | 9.04% | 5142.6 | 5.19% | 1.74 |
| Fra2 | GGATGACTCATC | -1.23E+03 | 12561 | 12.53% | 7906.4 | 7.98% | 1.57 |
| Fra1 | NNATGASTCATH | -1.22E+03 | 14460 | 14.42% | 9456.6 | 9.54% | 1.51 |
| JunB | RATGASTCAT | -1.18E+03 | 14270 | 14.23% | 9365.6 | 9.45% | 1.51 |
| Jun-AP1 | GATGASTCATCN | -1.13E+03 | 6857 | 6.84% | 3663.4 | 3.70% | 1.85 |
| PU.1:IRF8 | GGAAGTGAAAST | -9.88E+02 | 7023 | 7.00% | 3949.9 | 3.98% | 1.76 |
| SpiB | AAAGRGGAAGTG | -6.75E+02 | 7805 | 7.78% | 5016.1 | 5.06% | 1.54 |
| Bach2 | TGCTGAGTCA | -6.28E+02 | 5047 | 5.03% | 2933.6 | 2.96% | 1.70 |

| | | | | | | | |
|---|---|---|---|---|---|---|---|
| NF-E2 | GATGACTCAGCA | -2.94E+02 | 1548 | 1.54% | 777.5 | 0.78% | 1.97 |
| Bach1 | AWWNTGCTGAGTCAT | -2.84E+02 | 1471 | 1.47% | 734.7 | 0.74% | 1.99 |
| Nrf2 | HTGCTGAGTCAT | -2.33E+02 | 1298 | 1.29% | 666.7 | 0.67% | 1.93 |
| NFE2L2 | AWWWTGCTGAGTCAT | -2.18E+02 | 1336 | 1.33% | 709.4 | 0.72% | 1.85 |
| CTCF-SatelliteElement | TGCAGTTCCMVNWRTGGCCA | -1.70E+02 | 563 | 0.56% | 231.3 | 0.23% | 2.43 |
| Ronin | RACTACAACTCCCAGVAKGC | -1.62E+02 | 1239 | 1.24% | 707.4 | 0.71% | 1.75 |
| NFkB-p65-Rel | GGAAATTCCC | -1.32E+02 | 1675 | 1.67% | 1094 | 1.10% | 1.52 |
| T1ISRE | ACTTTCGTTTCT | -1.20E+02 | 483 | 0.48% | 218.9 | 0.22% | 2.18 |

**Supplementary Table 2. Motif enrichment for regions with increased accessibility in the SCADS higher-scoring group.**

A total of 75 transcription-factor binding motifs passed the significance threshold ($q < 0.05$) in HOMER. The table here listed those with top enrichment (ratio between % target sequence with motif and % background sequence with motif > 1.5). The table is sorted in ascending order of p-values. All motifs in this table have *q* value < 0.0001.

| Motif Name | Consensus | log p-value | # of Target Sequences with Motif(of 100297) | % of Target Sequences with Motif | # of Background Sequences with Motif(of 99045) | % of Background Sequences with Motif | Enrichment |
|---|---|---|---|---|---|---|---|
| RUNX1 | AAACCACARM | -1.25E+02 | 311 | 53.44% | 11296 | 23.39% | 2.28 |
| Etv2 | NNAYTTCCTGHN | -1.15E+02 | 266 | 45.70% | 8952.7 | 18.53% | 2.47 |
| Fli1 | NRYTTCCGGH | -1.13E+02 | 275 | 47.25% | 9591.1 | 19.86% | 2.38 |
| ETS1 | ACAGGAAGTG | -1.07E+02 | 270 | 46.39% | 9577.8 | 19.83% | 2.34 |
| RUNX | SAAACCACAG | -9.81E+01 | 231 | 39.69% | 7661.7 | 15.86% | 2.50 |
| ERG | ACAGGAAGTG | -9.72E+01 | 344 | 59.11% | 15183.2 | 31.43% | 1.88 |
| RUNX2 | NWAACCACADNN | -9.61E+01 | 261 | 44.85% | 9574.9 | 19.82% | 2.26 |
| GABPA | RACCGGAAGT | -8.81E+01 | 225 | 38.66% | 7789.7 | 16.13% | 2.40 |
| ETV1 | AACCGGAAGT | -8.69E+01 | 296 | 50.86% | 12410.2 | 25.69% | 1.98 |
| ETV4 | ACCGGAAGTG | -8.62E+01 | 248 | 42.61% | 9289.4 | 19.23% | 2.22 |
| EWS:ERG-fusion | ATTTCCTGTN | -8.26E+01 | 222 | 38.14% | 7890.9 | 16.34% | 2.33 |
| RUNX-AML | GCTGTGGTTW | -8.12E+01 | 225 | 38.66% | 8144.3 | 16.86% | 2.29 |
| Ets1-distal | MACAGGAAGT | -7.88E+01 | 121 | 20.79% | 2764.4 | 5.72% | 3.63 |

| | | | | | | | |
|---|---|---|---|---|---|---|---|
| EWS:FLI1-fusion | VACAGGAAAT | -7.38E+01 | 170 | 29.21% | 5354.1 | 11.08% | 2.64 |
| Elk1 | HACTTCCGGY | -6.28E+01 | 133 | 22.85% | 3889.9 | 8.05% | 2.84 |
| ETS:RUNX | RCAGGATGTGGT | -6.11E+01 | 60 | 10.31% | 854.6 | 1.77% | 5.82 |
| PU.1 | AGAGGAAGTG | -5.74E+01 | 146 | 25.09% | 4818.1 | 9.97% | 2.52 |
| ETS Promoter | AACCGGAAGT | -5.28E+01 | 93 | 15.98% | 2343.7 | 4.85% | 3.29 |
| EHF | AVCAGGAAGT | -4.69E+01 | 260 | 44.67% | 12820.1 | 26.54% | 1.68 |
| Elf4 | ACTTCCKGKT | -4.64E+01 | 210 | 36.08% | 9407 | 19.48% | 1.85 |
| ELF1 | AVCCGGAAGT | -4.39E+01 | 112 | 19.24% | 3652.8 | 7.56% | 2.54 |
| Elk4 | NRYTTCCGGY | -4.27E+01 | 115 | 19.76% | 3871.5 | 8.02% | 2.46 |
| ELF3 | ANCAGGAAGT | -3.53E+01 | 167 | 28.69% | 7472.9 | 15.47% | 1.85 |
| IRF2 | GAAASYGAAASY | -2.62E+01 | 47 | 8.08% | 1205.7 | 2.50% | 3.23 |
| ELF5 | ACVAGGAAGT | -2.03E+01 | 144 | 24.74% | 7349.4 | 15.22% | 1.63 |
| IRF4 | ACTGAAACCA | -2.01E+01 | 111 | 19.07% | 5187.5 | 10.74% | 1.78 |
| SPDEF | ASWTCCTGBT | -1.93E+01 | 175 | 30.07% | 9628.8 | 19.93% | 1.51 |
| IRF8 | GRAASTGAAAST | -1.51E+01 | 82 | 14.09% | 3803.5 | 7.87% | 1.79 |
| IRF1 | GAAAGTGAAAGT | -1.51E+01 | 48 | 8.25% | 1776.1 | 3.68% | 2.24 |

**Supplementary Table 3. Motif enrichment for regions with decreased accessibility in the SCADS higher-scoring group.**

A total of 3 transcription-factor binding motifs passed the significance threshold (q < 0.1) in HOMER. The table here listed those with top enrichment (ratio between % target sequence with motif and % background sequence with motif > 1.5). The table is sorted in ascending order of p-values. All motifs in this table have *q* value < 0.1.

| Motif Name | Consensus | log *p*-value | # of Target Sequences with Motif (of 100297) | % of Target Sequences with Motif | # of Background Sequences with Motif (of 99045) | % of Background Sequences with Motif | Enrichment |
|---|---|---|---|---|---|---|---|
| CTCF | AYAGTGCCMYCTRGTGGCCA | -1.18E+02 | 72 | 34.29% | 1635.7 | 3.29% | 10.42 |
| BORIS | CNNBRGCGCCCCCTGSTGGC | -7.50E+01 | 83 | 39.52% | 4384.4 | 8.83% | 4.48 |
| Bach1 | AWWNTGCTGAGTCAT | -7.61E+00 | 7 | 3.33% | 322.1 | 0.65% | 5.12 |

**Supplementary Table 4. Correlation between chromatin accessibility and disease score for $CD8^+$ CM T cells**

The table here is a list of the correlations between chromatin accessibility and IBD SCADS scores at 42 variant loci sorted by descending order of absolute correlation. P-values were adjusted using the Benjamini-Hochberg procedure (FDR). We categorized the variants into three groups based on their accessibility correlation with the disease score: positive (Pearson r > 0.2), negative (Pearson r < -0.2) and no correlation (the rest).

| SNP | Correlation | *p* adjusted | Category |
|---|---|---|---|
| rs67289879 | 0.992 | 0.035 | Positive |
| rs7933433* | 0.964 | 0.120 | Positive |
| rs194746 | 0.953 | 0.120 | Positive |
| rs17056705 | 0.944 | 0.120 | Positive |
| rs259958 | 0.937 | 0.120 | Positive |
| rs6677188 | 0.934 | 0.120 | Positive |
| rs185034120 | 0.901 | 0.161 | Positive |
| rs12922863 | 0.898 | 0.161 | Positive |
| rs72850698 | 0.870 | 0.194 | Positive |
| rs3181374 | 0.866 | 0.194 | Positive |
| rs1551398 | 0.855 | 0.194 | Positive |
| rs7524424 | 0.844 | 0.197 | Positive |
| rs744166 | 0.830 | 0.197 | Positive |
| rs28510097 | 0.829 | 0.197 | Positive |
| rs7240004 | 0.820 | 0.197 | Positive |
| rs3024505 | 0.776 | 0.246 | Positive |
| rs77239361 | 0.756 | 0.267 | Positive |
| rs72870519 | 0.731 | 0.293 | Positive |
| rs78703675 | 0.634 | 0.438 | Positive |
| rs71381201 | 0.614 | 0.449 | Positive |
| rs75944240 | 0.607 | 0.449 | Positive |
| rs4303859 | 0.591 | 0.458 | Positive |
| rs3743427 | 0.485 | 0.606 | Positive |
| rs2057657 | 0.475 | 0.606 | Positive |

| | | | |
|---|---|---|---|
| rs112401631 | 0.399 | 0.662 | Positive |
| rs149290349 | 0.390 | 0.662 | Positive |
| rs140020875 | 0.373 | 0.662 | Positive |
| rs633059 | 0.300 | 0.749 | Positive |
| rs9834996 | -0.939 | 0.120 | Negative |
| rs3746703 | -0.904 | 0.161 | Negative |
| rs12097268 | -0.859 | 0.194 | Negative |
| rs8122494 | -0.822 | 0.197 | Negative |
| rs77980086 | -0.806 | 0.210 | Negative |
| rs138504748 | -0.431 | 0.657 | Negative |
| rs112694524 | -0.376 | 0.662 | Negative |
| rs56235845 | 0.164 | 0.899 | None |
| rs137965 | 0.093 | 0.975 | None |
| rs11677002 | 0.062 | 0.976 | None |
| rs12132298 | 0.037 | 0.976 | None |
| rs12132349 | 0.037 | 0.976 | None |
| rs153146 | -0.007 | 0.991 | None |
| rs11645239 | -0.230 | 0.828 | None |

* Note: Highlighted in results

**Supplementary Table 5. Enrichment of immune cell-specific regulons**
Enrichment analysis results using Fisher's test for our SNP associated target genes in established regulons is provided here. The table is sorted by descending order of enrichment.

| TF Regulon | Regulon Size | Overlap genes | Enrichment | *p* adjusted | Regulon activity in CD8 T cells |
|---|---|---|---|---|---|
| TCF7_(+)* | 18 | *ZFP36L2, ETS1* | 65.122 | 0.075 | 0.199 |
| DDIT3_(+) | 281 | *TRIB1, PTGER4, ZFP36L1, ERRFI1* | 8.768 | 0.106 | 0.162 |
| EGR2_(+) | 235 | *TRIB1, ZFP36L1, IRF1* | 7.644 | 0.193 | 0.062 |
| SP1_(+) | 500 | *PRKCB, IRF1, LPP, TBC1D1, ERRFI1* | 6.289 | 0.106 | 0.064 |
| KLF9_(+) | 599 | *ZFP36L2, IL10, TBC1D1, CTIF, STAT3* | 5.223 | 0.128 | 0.124 |
| CREM_(+) | 1087 | *ZFP36L2, CIITA, PTGER4, IRF1, TNFSF8, IL23R, PTPN2, ERRFI1* | 4.971 | 0.075 | 0.105 |
| ETS2_(+)* | 1608 | *ZFP36L2, CIITA, TRIB1, ZFP36L1, IRF1, IL10, TNFSF8, CTIF, PTPN2, TPD52L2* | 4.399 | 0.075 | 0.078 |
| RELB_(+) | 1604 | *SMARCE1, ZFP36L1, IRF1, IL10, TNFSF8, CCND3, PTPN2, STAT3, IP6K2* | 3.822 | 0.106 | 0.083 |
| MAFF_(+) | 1240 | *ZFP36L2, TRIB1, IRF1, IL10, CTIF, IL19, TPD52L2* | 3.651 | 0.162 | 0.091 |
| HIF1A_(+) | 1725 | *TRIB1, ZFP36L1, IRF1, IL10, CTIF, PTPN2, STAT3, ERRFI1, TPD52L2* | 3.531 | 0.122 | 0.089 |
| BACH1_(+) | 1815 | *PRKCB, TRIB1, ZFP36L1, IRF1, TNFSF8, LPP, TBC1D1, ERRFI1, TPD52L2* | 3.339 | 0.128 | 0.075 |
| ATF3_(+) | 1350 | *ZFP36L2, TRIB1, PTGER4, ZFP36L1, IRF1, PTPN2, STAT3* | 3.334 | 0.193 | 0.117 |
| FOSB_(+) | 1618 | *ZFP36L2, TRIB1, PTGER4, ZFP36L1, IRF1, TNFSF8, LPP, IL23R* | 3.246 | 0.180 | 0.117 |
| FOS_(+) | 2972 | *ZFP36L2, CIITA, TRIB1, PTGER4, ZFP36L1, IRF1, ZNF831, IL10, LPP, CCND3, CTIF, PTPN2, STAT3* | 3.238 | 0.106 | 0.103 |

| | | | | | |
|---|---|---|---|---|---|
| FOSL2_(+) | 2182 | *ZFP36L2, PRKCB, TRIB1, PTGER4, ZFP36L1, IRF1, IL10, ERRFI1, RORC, TPD52L2* | 3.140 | 0.128 | 0.086 |
| IRF1_(+) | 3189 | *ZFP36L2, PRKCB, POM121C, TRIB1, PTGER4, IRF1, IL10, LPP, TBC1D1, STAT3, ERRFI1, RORC, IP6K2* | 2.979 | 0.122 | 0.079 |
| REL_(+) | 2099 | *ZFP36L2, PRKCB, CIITA, POM121C, PTGER4, ZFP36L1, IRF1, LPP, TBC1D1* | 2.843 | 0.198 | 0.072 |
| SPIB_(+) | 2440 | *SMARCE1, PRKCB, CIITA, UCKL1, PTGER4, ZFP36L1, IRF1, TBC1D1, CCND3, IP6K2* | 2.768 | 0.193 | 0.101 |
| ELF1_(+) | 3801 | *SMARCE1, ZFP36L2, POM121C, PTGER4, IRF1, ZNF831, TNFSF8, LPP, IL23R, PTPN2, STAT3, IL19, ETS1, IP6K2* | 2.712 | 0.128 | 0.085 |
| ETS1_(+) | 6521 | *SMARCE1, ZFP36L2, PRKCB, UCKL1, POM121C, PTGER4, IRF1, ZNF831, TNFSF8, NELFCD, LPP, CCND3, RMI2, STAT3, ERRFI1, ELP6, RORC, ETS1, IP6K2* | 2.310 | 0.193 | 0.080 |

* Note: Highlighted in results

**Supplementary Table 6. Correlation between chromatin accessibility and disease score for colon epithelial cells**

The table here is a list of the correlations between chromatin accessibility at 24 variant loci and IBD SCADS scores. They are sorted by descending order of absolute correlation within each category: Positive (Pearson r > 0.2), negative (Pearson r < -0.2) and no correlation (the rest). P-values were adjusted using the Benjamini-Hochberg procedure (FDR).

| SNP | Correlation | *p* adjusted | Category |
|---|---|---|---|
| rs2816972 | 0.993 | 0.011 | Positive |
| rs138504748 | 0.947 | 0.058 | Positive |
| rs35260072 | 0.933 | 0.070 | Positive |
| rs194746 | 0.906 | 0.103 | Positive |
| rs2284553 | 0.877 | 0.114 | Positive |
| rs9486287 | 0.876 | 0.114 | Positive |
| rs2143607 | 0.658 | 0.321 | Positive |
| rs4807569 | 0.633 | 0.336 | Positive |
| rs7553638* | -0.990 | 0.011 | Negative |
| rs56235845 | -0.989 | 0.011 | Negative |
| rs1317209* | -0.978 | 0.023 | Negative |
| rs12654812 | -0.954 | 0.056 | Negative |
| rs3746703 | -0.875 | 0.114 | Negative |
| rs77980086 | -0.843 | 0.137 | Negative |
| rs73205554 | -0.841 | 0.137 | Negative |
| rs116886623 | -0.792 | 0.184 | Negative |
| rs11645239 | -0.786 | 0.184 | Negative |
| rs140020875 | -0.725 | 0.249 | Negative |
| rs73432869 | -0.528 | 0.456 | Negative |
| rs149290349 | -0.504 | 0.465 | Negative |
| rs3743427 | -0.445 | 0.517 | Negative |
| rs16940186 | 0.082 | 0.961 | None |
| rs3820328 | 0.061 | 0.961 | None |
| rs11245346 | 0.031 | 0.961 | None |

* Note: Highlighted in results

1. Granja, J. M. *et al.* Single-cell multiomic analysis identifies regulatory programs in mixed-phenotype acute leukemia. *Nat. Biotechnol.* **37**, 1458–1465 (2019).
2. Trapnell, C. *et al.* The dynamics and regulators of cell fate decisions are revealed by pseudotemporal ordering of single cells. *Nat. Biotechnol.* **32**, 381–386 (2014).
3. Cao, J. *et al.* The single-cell transcriptional landscape of mammalian organogenesis. *Nature* **566**, 496–502 (2019).
4. Csárdi, G. *et al. Igraph for R: R Interface of the Igraph Library for Graph Theory and Network Analysis*. (Zenodo, 2026). doi:10.5281/ZENODO.7682609.
